# Structural and cellular transcriptome foundations of human brain disease

**DOI:** 10.1101/2021.05.12.443911

**Authors:** Yashar Zeighami, Trygve E. Bakken, Thomas Nickl-Jockschat, Zeru Peterson, Anil G. Jegga, Jeremy A. Miller, Alan C. Evans, Ed S. Lein, Michael Hawrylycz

## Abstract

Genes associated with risk for brain disease exhibit characteristic expression patterns that reflect both anatomical and cell type relationships. Brain-wide transcriptomic patterns of disease risk genes provide a molecular based signature for identifying disease association, often differing from common phenotypic classification. Analysis of adult brain-wide transcriptomic patterns associated with 40 human brain diseases identified five major transcriptional patterns, represented by tumor-related, neurodegenerative, psychiatric and substance abuse, and two mixed groups of diseases. Brain disease risk genes exhibit unique anatomic transcriptomic signatures, based on differential co-expression, that often uniquely identify the disease. For cortical expressing diseases, single nucleus data in the middle temporal gyrus reveals cell type expression gradients separating neurodegenerative, psychiatric, and substance abuse diseases. By homology mapping of cell types across mouse and human, transcriptomic disease signatures are found largely conserved, but with psychiatric and substance abuse related diseases showing important specific species differences. These results describe the structural and cellular transcriptomic landscape of disease in the adult brain, highlighting significant homology with the mouse yet indicating where human data is needed to further refine our understanding of disease-associated genes.

## Introduction

Brain diseases are increasingly recognized as major causes of death and disability worldwide (1–3). These diverse and multifactorial diseases may be largely grouped into cerebrovascular, neurodegenerative, movement related, psychiatric disorders, developmental and congenital disorders, substance abuse disorders, brain tumors, and a set of other brain-related diseases (*Institute for Health Metrics* (IHME), healthdata.org). The etiology of brain-related diseases and their genetics is complex and widely studied (4–6). However, phenotypic classification of brain diseases is challenging and does not uniquely partition characteristics of genetic risk, disease manifestation, and treatment. Except for Mendelian diseases arising from single gene mutations, most brain disorders present as a complex interplay between genetics and environment through interaction of the brain transcriptome and its regulatory network. Genetic analysis of brain disease, through profiling of tissues, cells, and more recently at the resolution of single nuclei (7) addresses this complexity providing a means for population scale sampling to disentangle basic molecular relationships (8, 9). The economic impact of brain diseases also varies substantially, as reflected in the comprehensive and annually updated *Global Burden of Disease Study* (10) (**Suppl. Figure 1**).

Investigating the neuroanatomy of major transcriptomic relationships for brain diseases and their relationship to cell type provides a novel means of disease comparison and classification, indicating directions and potential for follow up as brain-wide cellular data becomes available. The premise of the present study is based on the hypothesis that spatial and temporal co-expression of disease genes is indicative of a potential interaction between these genes (11, 12). Studying brain samples from donor populations exhibiting coherent transcriptomic and anatomic relationships of disease-related genes, both in neurotypical and diseased brains and at multiple scales, promises important insight in developing further approaches to study the pathophysiology of brain disorders. Large scale transcriptome profiling of the human brain has already produced useful resources for exploring the genetics of neurotypical and disease states (13–16) and for describing the larger scale relationship of brain diseases and the neuroanatomy of transcriptomic patterning (13, 17).

Transcriptomic relationships at a mesoscale, intermediate between the larger brain structures (e.g., cortex, hypothalamus) and those at cellular resolution, provide a natural framework and starting point for classifying broad disease associations in comparison with common phenotypic grouping. Starting with the *Allen Human Brain Atlas* (human.brain-map.org), (13, 14), we investigated anatomic patterning and differential expression of the transcriptional patterns of genes for 40 brain-related disorders across 104 structures from cortex, hippocampus, amygdala, basal ganglia, epithalamus, thalamus, ventral thalamus, hypothalamus, mesencephalon, cerebellum, pons, pontine nuclei, myelencephalon, ventricles, and white matter. Using single nucleus data from the human middle temporal gyrus, we subsequently characterize a subset of 24 diseases with primary expression in cortex by comparing expression of cell types from a taxonomy of 45 inhibitory, 24 excitatory, 6 non-neuronal types, with special attention to psychiatric diseases. This multiresolution approach combining tissue based and single nucleus data connects mesoscale anatomic analysis with cell types of the cortex and is a recognized approach for extracting information from tissue-based sampling (18, 19). Finally, juxtaposing these results with single cell data in mouse (celltypes.brain-map.org) (15, 20) allows identification of human specific potential cell type differences.

### Brain disorders and associated genes

The diseases selected are representative of seven phenotypic classes from the *Global Burden of Disease Study* (referred to as ***GBD*** classes in this study). The important group of cerebrovascular diseases were excluded due to limitations of representative endothelial and pericyte cell types and related blood cells in data sources. To identify gene-disease associations, we used the *DisGeNET* database (www.disgenet.org) (21–23) a platform aggregated from multiple sources including curated repositories, GWAS catalogs, animal models and the scientific literature. From an initial survey of the *Online Mendelian Inheritance in Man (OMIM)* (www.omim.org) repository we first identified 549 brain-related diseases (14) which were intersected with the *DisGeNET* repository. We required reported gene-disease associations to be present in at least one confirmed curated source (See https://www.disgenet.org/dbinfo), and with a minimum of 10 genes per disease. For each disease, the main variant of the disease was selected with rare familial and genetic forms not included. This conservative selection resulted in 40 major brain disorders with 1646 unique associated genes. **Suppl. Table 1** contains definitions, gene sets, and metadata identifying each disease. (**Methods**).

Gene sets associated with brain disease vary widely in size and the proportion of shared genes between diseases is known to be correlated with phenotypic similarity (*ρ* = 0.40, *p* = 6.0 × 10^−3^), based on clinical manifestations (24, 25). Gene set sizes range widely from, frontotemporal lobar degeneration (g=11), to schizophrenia (g=733) and distribute across GBD classes as (number, % unique to GBD class) psychiatric (1107, 0.723), neurodegenerative (257, 0.513), substance abuse (212, 0.320), brain tumors (168, 0.667), developmental disorders (139, 0.676), movement related (136, 0.272) and other brain-related (123, 0.414). The comorbidity of psychiatric diseases and substance abuse is well established (26) and **Supp. Fig. 2A** shows the largest gene set intersection (g=132) between GBD classes psychiatric and substance abuse, with 62% of substance abuse genes also associated with psychiatric disorders. Movement disorders are also commonly found in neurodegenerative diseases (27), with GBD neurodegenerative sharing 30% (g=41) of movement related genes, while GBD tumor based and developmental sharing the least with other classes (2.5%, 2.6% respectively). Clustering the 40 diseases and disorders based on relative pairwise gene set intersection (Jaccard) shows moderate agreement with GBD phenotypic groupings (**Supp. Fig. 2B),** with the highest percentage of shared genes among psychiatric disorders 7.64% (p=1.55 × 10^-4^), followed by substance abuse 6.33% (p=2.82 × 10^-4^), and brain tumors 5.43% (p=8.35 × 10^-3^). (Significance is likelihood of observed percentage corrected for GBD class size.) Functional enrichment analysis (https://toppgene.cchmc.org) both of GBD pooled genes, and genes unique to individual diseases, describe known annotations (**Suppl. Fig.3**, **Suppl. Table 2**.)

### Structural transcriptomic profile of brain diseases

Expression profiles from the *Allen Human Brain Atlas* (AHBA, https://human.brain-map.org) from 6 neurotypical donor brains are used to summarize major neuroanatomical relationships of genes associated with the 40 diseases. Using an ontology of 104 structures (**Suppl. Table 3**) including cortex (CTX, consisting of 8 structures), hippocampus (HIP,7), amygdala (AMG,6), basal ganglia (BG,12), epithalamus (ET,3), thalamus (TH,12), hypothalamus (HY,16), mesencephalon (MES, 11), cerebellum and cerebellar nuclei (CB,4), pons and pontine nuclei (P,10) myelencephalon (MY, 12), ventricles (V,1), white matter (WM,2), we obtained a mean transcriptomic disease profile by averaging expression for genes associated with each of 40 diseases across the 104 structures and z-score normalizing (**Figure 1, Suppl. Table 4)**. Performing hierarchical clustering with Ward linkage using Pearson correlation (**Methods**) presents brain wide transcriptomic associations in five primary *Anatomic Disease Groups* (**ADG 1-ADG 5**) interpretable with respect to GBD classification (**Fig. 1A**, left color bar) as tumor related (**ADG 1**), neurodegenerative (**ADG 2**), psychiatric, substance abuse, and movement disorders (**ADG 3**), and two mixed groups *without* developmental, psychiatric or tumor diseases (**ADG 4,5**). We obtain an anatomic representation of transcriptomic patterning by averaging gene expression within each ADG group across the 104 brain structures (**Fig. 1**), whose major anatomy is described as **ADG 1:** thalamus, brain stem, ventricle wall, white matter, **ADG 2:** cortico-thalamic, brain stem, white matter, **ADG 3:** (telencephalon) cortex, thalamus, hippocampus, amygdala, basal ganglia, **ADG 4:** basal ganglia, hypothalamus, brain stem, and **ADG 5:** thalamus, hypothalamus, brain stem (**Suppl. Fig. *4***). ADG transcriptome signatures are consistent across subjects as individual brain holdout analysis (**Supp. Figs. 5,6**, **Methods**) finds that both the correlation of expression across structures and differential relationships between ADG groups at a fixed structure are preserved within the AHBA subjects, indicating reproducible signatures across subjects.

**Figure 1.**
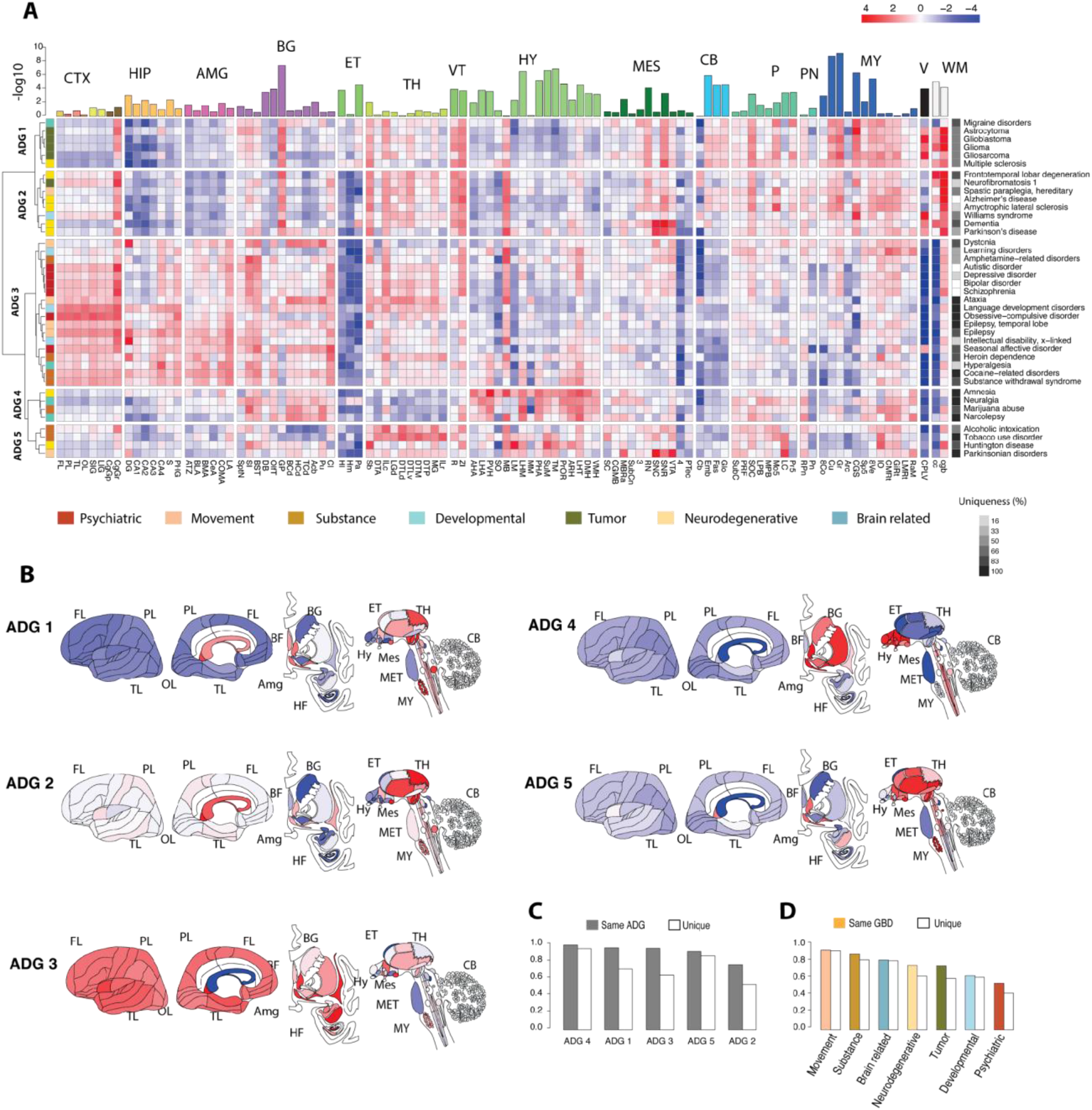
Transcriptome patterning of major brain diseases. A) Mean gene expression profiles for genes associated with 40 major brain diseases and disorders profiled over 104 anatomic structures (**Suppl. Table 3**) from 15 major regions cortex (CTX), hippocampus (HIP), amygdala (AMG), basal ganglia (BG), epithalamus (ET), thalamus (TH), ventral thalamus (VT), hypothalamus (HY), mesencephalon (MES), cerebellum (CB), pons (P), pontine nuclei (PN), myelencephalon (MY), ventricles (V), white matter (WM) and z-score normalized. Hierarchical clustering based on mean profile and z-score normalized yields five primary anatomic disease groups **ADG 1 – ADG 5**. Row annotation (left bar) shows phenotypic GBD classification is variable with ADG membership. Column bar annotation is five group (ADG) ANOVA for expression variability at a fixed structure. Row annotation (right bar), correlation-based uniqueness measure of disease signature across AHBA brains; 100% means transcriptomic profile uniquely identifies disease in all subjects (**Methods**.) B) Adult brain graphic showing anatomic patterning of classes ADG 1-5, C) Reproducibility of ADG profiles. Left: (Solid) Frequency across subjects that the transcriptomic signature is most closely correlated with a signature from the same ADG in other subjects. (Open) Frequency that exact disease is identified in other subjects. Right: Similar analysis for diseases by GBD groups showing movement and substance abuse as the most identifiable disease groups.

To quantify the most significant expression differences between ADG groups, we apply ANOVA for mean differences in expression across ADG groups at each structure (BH corrected p-values, top panel, **Fig. 1**.) Particularly striking in **Fig. 1A** is the white matter signature common to tumor and neurodegenerative diseases (**ADG 1-2**), effectively absent in psychiatric disorders and diseases of addiction (**ADG 3-4**) (28), and substantial enrichment of telencephalic expression (CTX, HIP, AMG, BG) in **ADG 3** (29, 30). The most significant transcriptomic variation in disease genes occurs across the diverse nuclei of lower brain structures: in the hypothalamus (e.g., cuneate nucleus (Cu, p < 3.35 × 10^-8^), tuberomammillary nucleus (TM, 1.30 × 10^-6^), supramammillary nucleus (SuM, 2.07 × 10^-6^)), in the myelencephalon (gracile nucleus (GR, *p* < 1.48 × 10^-8^)), central glial substance (CGS, 3.86 × 10^-6^), in the basal ganglia (globus pallidus (GP, 5.01 × 10 ^−7^)), and cerebellar nuclei (CN, and white matter, in particular, corpus callosum (cc, 5.42 × 10^-5^). Global variation across disease genes is not significant in the thalamus (TH, 0.338), myelencephalon (0.247), and cerebellum (CB, 0.966), whereas differences in telencephalic expression between psychiatric, substance abuse, and movement groups (**ADG 3**) and other ADGs is demonstrated by applying paired t-tests between groups (**Suppl. Fig. 7**). Here **ADG 1** and **ADG 3** are distinguished through differences in frontal lobe (FL, *p* < 2.71 × 10^-7^), hippocampus, dentate gyrus (DG, 3.46 × 10^-6^), and amygdala, basomedial nucleus (BMA, 4.49 × 10^-10^). The distinction between **ADG 1** and **ADG 2** is more subtle with variation in cortex (frontal lobe, FL, 2.82 × 10^-3^), epithalamus (lateral habenular nucleus, (HI, 2.85 × 10^-5^), and mesencephalon (pretectal region, (PTec, 6.24 × 10^-5^). Finally, **ADG 4**, **5** differences are characterized by diencephalon expression: thalamus, anterior group of nuclei (DTA, 3.01 × 10^-7^), lateral group of nuclei, dorsal division, (DTLv, 6.47 × 10 ^−9^), and hypothalamus, posterior hypothalamic area (PHA, 1.21 × 10^-6^).

The complex anatomic organization of gene expression reflected in **Fig. 1** associates diseases with common phenotypic classification by the GBD study but with important divergences (**Fig. 1a,** left sidebar). **ADG 1** and **ADG 2,** driven by co-expression in the diencephalon, myelencephalon and white matter, comprise tumor based and neurodegenerative diseases with the addition of migraine disorders and multiple sclerosis within **ADG 1** and Williams syndrome and hereditary spastic paraplegia within **ADG 2**. The common association of all psychiatric diseases, and most movement, and substance disorders in **ADG 3** is driven by strong telencephalic patterning, while **ADG 4** and **5** comprise diseases from mixed phenotypic classes not well associated with other ADGs. These latter groups are highly mixed phenotypic groups associating amnesia with neuralgia, and Parkinsonian disorders with substance abuse. Although ADGs reflect the broadest transcriptome patterning there is important variation within groups. For example, substantia nigra (SNC, SNR) and ventral tegmental area (VTA) enrichment is seen in genes associated with dementia, Parkinson’s and related disorders (31), strong expression in the thalamus is identified for alcohol and tobacco use disorders (32), and genes associated with amnesia exhibit hypothalamic expression atypical of other neurodegenerative disease except for the mammillary body (MB). **Fig. 1** illustrates the rich anatomic structure of disease gene expression and remarkably, the division and structure of **ADG** groups is largely preserved upon *removing* genes common between pairs of diseases (**Suppl. Fig. 8**) showing that major ADG groups are driven by distinct co-expressing genes. Ten additional diseases with limited gene sets (5–10) are presented in **Suppl. Fig. 9**, with their relationship to ADG groups.

While the expression of disease genes may vary considerably in a population (33, 34), the anatomic expression signature of each disease in an individual brain is most closely correlated with a disease in the same ADG group in other brains (**ADG 1-5**: 96.7, 77.0, 96.1, 100.0, 92.5 %), and typically identifies the exact disease in other subjects (**Fig. 1C, Methods**). In particular, the expression pattern associated with the **ADG 3** group diseases ataxia, language development disorders, temporal lobe epilepsy, obsessive compulsive disorder, and cocaine-related disorder most closely correlates with these same diseases in each of the subjects. Similarly, in **ADG 4, 5,** genes associated with Parkinsonian disorders, Huntington’s disease, amnesia, narcolepsy, neuralgia, and tobacco use disorder exhibit highly unique profiles due to consistent, complex and differentiated expression in the basal ganglia, hypothalamus, and myelencephalon. Conversely, the mesoscale transcriptomic profile of **ADG 2** Alzheimer’s disease and amyotrophic lateral sclerosis, and **ADG 3** bipolar disorder, autistic disorder, and schizophrenia are less unique to those diseases. Phenotypically, GBD movement disorders and substance abuse have the most consistent anatomic signatures (94.0,89.5%) (**Fig. 1D**), while psychiatric and developmental diseases the least (64.0%, 55.0%). The ability to uniquely identify a disease from its anatomic signature indicates a finer transcriptomic patterning and is a bridge to cell type analysis.

### Reproducible transcription patterns of brain diseases

The neuroanatomy of transcription patterns for disease risk genes can be further studied by identifying conserved differential expression relationships, which provides a bridge to implicated cell types. *Differential stability* (DS), introduced in (13), is quantified as the mean Pearson correlation ρ of expression between pairs of specimens over a fixed set of anatomic regions, and measures the fraction of preserved differential relationships between anatomic regions for a set of subjects. For example, the gene *GRIA2* with high DS (ρ = 0.918), (**Figure 2A**) is implicated in bipolar disorder (35), schizophrenia (36), and substance withdrawal syndrome (37) and has a highly reproducible brain wide expression profile across AHBA subjects with highest expression in hippocampus and amygdala.

**Figure 2.**
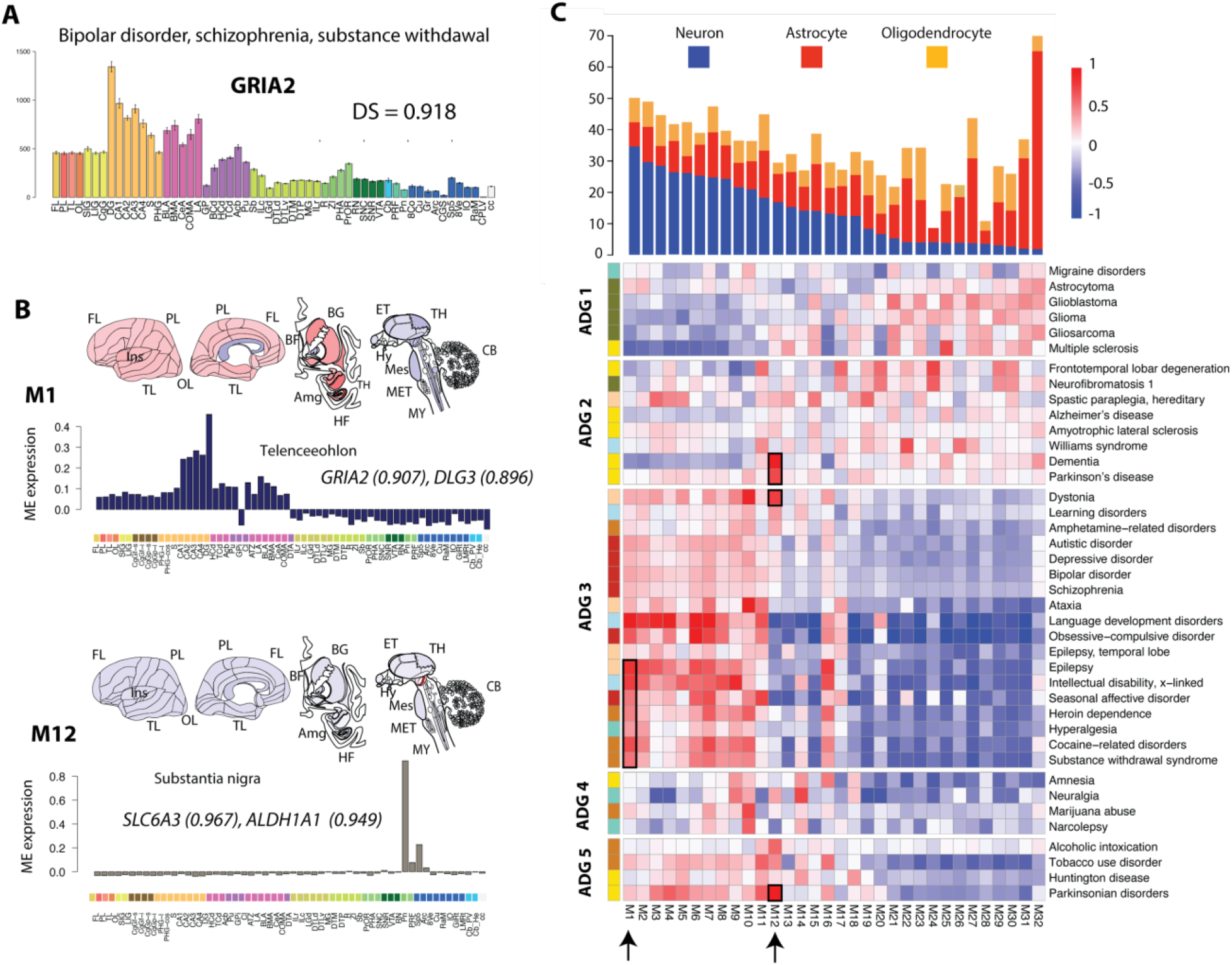
Reproducible transcription patterns in human brain diseases. A) Expression profile for gene GRIA2 with error bars shown over 56 structures (**Suppl. Table 3**, human.brain-map.org.) Differential stability (DS) measures reproducible expression patterns from the Allen Human Brain Atlas (AHBA) (13). B) Canonical eigengenes for **M1** *telencephalic* (language development, epilepsy) and **M12** *substantia nigra* (Parkinson’s disease, dementia), with module correlation for representative genes. C) Map of canonical expression modules **M1-M32** mapping anatomic co-expression to diseases. Disease genes are correlated with each module independently and normalized (**Methods**), disease ordering is the same as in **Fig. 1**. Modules **M1**-**M32** are ordered based on their neuronal, astrocyte, oligodendrocyte cell type content derived in (13). Arrows and boxes indicate **M1** and **M12**. Other disease representative modules described in **Suppl. Fig 12**.

Although disease genes show a marginally significant (p<0.031) difference in their expression levels compared to non-disease associated (**Suppl. Fig. 10A**), these genes have a significantly higher percentage (**Suppl. Fig. 10B**) of differentially stable genes, particularly for substance abuse, (mean DS=0.702, p<4.7 × 10^-21^), psychiatric (DS=0.675, p<3.17 × 10^-17^), and movement disorders (DS=0.635, p<9.1× 10^-17^). Notably, the stability of **ADG 3** (median 0.625, p<4.11 × 10^-70^), **ADG 4** (0.642, 1.24 × 10^-6^) and **ADG 5** (0.644, 2.95 × 10^-14^) genes are markedly higher than **ADG 1** (0.592, 1.02 × 10^-6^) and **ADG 2** (0.582, 7.70 × 10 ^−7^) indicating a higher percentage of neuronal cell types and structural markers for diseases in these groups. **Suppl. Fig. 10C** shows the distribution of DS genes for each disease, confirming that diseases with higher DS are those with structurally reproducibly expression signatures (**Suppl. Fig. 11**). High DS disease genes are also substantially enriched for cell type processes, e.g., anterograde trans-synaptic signaling, (low DS 1.8 × 10^-4^, high 2.9 × 10^-12^), presynaptic membrane (low DS 0.049, high 4.62 × 10^−7^), indicating high DS to select for cell type specificity.

A characterization of the reproducible gene co-expression patterns (14) in the *Allen Human Brain Atlas* using the top half of DS genes (DS > 0.5284, g = 8,674) previously identified 32 primary transcriptional patterns, or *modules,* each represented by a characteristic expression pattern (i.e., eigengene) across brain structures and ordered by cell type content. **Fig. 2B** illustrates the membership of disease risk genes to modules for two representative modules **M1** and **M12**. Module **M1** has strong telencephalic expression in the hippocampus, in particular dentate gyrus, and representative genes include *GRIA2* (correlation to eigengene, ρ=0.907), and *DLG3* (ρ=0.896).

Alterations in glutamatergic neurotransmission have known associations with psychiatric and neurodevelopmental disorders and mutations in *GRIA2* have been related with these disorders (33–35). **M12** is a unique neuronal marker of substantia nigra *pars compacta*, *pars reticulata*, and ventral tegmental area and provides a clearer connection of dystonia, Parkinson’s disease, and dementia for these comorbidities (**Fig. 2C**). Both the dopamine transporter gene *SLC6A3* (ρ=0.967), a candidate risk gene for dopamine or other toxins in the dopamine neurons (38, 39) and aldehyde dehydrogenase-1 (*ALDH1A1,* ρ=0.949), whose polymorphisms are implicated in alcohol use disorders, map to module **M12** (ρ=0.949) (40). Brain wide association of expression module profiles may potentially implicate genes without previous association to a given disease, particularly when that profile is highly conserved between donors.

We map brain-related diseases to the canonical patterns by finding the closest correlated module eigengene for each disease gene (**Supp. Table 5**). **Figure 2C s**hows the normalized mean correlation of the 40 disease associated gene sets with the module **M1-M32** eigengenes ordered by **ADG 1-5** as in **Fig.1** (**Methods**). The basic cell class composition of neuronal, oligodendrocyte, astrocyte of AHBA tissue samples was determined from earlier single cell studies (13) and the modules **M1-M32** ordered by decreasing proportion of neuron-enriched cells. Interestingly, **Fig. 2C** clarifies the distinction between ADG groups of **Fig. 1,** shows major cell type content, and illustrates the anatomic co-expression patterns of brain diseases. Primarily tumor-based **ADG 1** map to modules **M21-M32** having enriched glial content (p<2.413× 10^−15^), while **ADG 3** psychiatric and substance abuse related diseases map to neuronal enriched patterned modules **M1-M10** (2.2× 10^-16^). By contrast the neurodegenerative disorders of **ADG 2** including Alzheimer’s, Parkinson’s, ALS, and frontotemporal lobe degeneration show more uniform distribution across the modules, and clearly separating this group from **ADG 1** (1.55× 10^−15^). **ADG 4,5** are both enriched in specific anatomic markers, e.g., **M10** (striatum), narcolepsy, marijuana, **M14** (hypothalamus), neuralgia, amnesia, **M11** (thalamus, Parkinsonian and tobacco use disorders), **M12** (substantia nigra, Parkinsonian and alcoholic intoxication) yet have lower expression in neuronal modules **M1-12** than **ADG 3** (1-sided, p<3.84× 10^-13^). The canonical module distribution of **Fig. 2C** validates the clustering of **Fig. 1**, improves the unique disease identification for ADG and provides a bridge to understanding cell type content (**Suppl. Figs. 13,14**).

### Disease genes and cell types of middle temporal gyrus

A primary telencephalic expression pattern is common to diseases of **ADG 3**, and while neural systems level analysis describes brain-wide anatomic relationships, it is limited in its ability to implicate specific cell types in diseases (12, 41). To more finely describe these diseases, we now restrict to those 24 diseases having *higher* than median cortical expression in the brain wide analysis (**Figs. 1–2**); essentially the entirety of **ADG 3** and several neurodegenerative diseases from **ADG 2**. We used human single nucleus (snRNA-seq) data from eight donor brains (15,928 nuclei) from the middle temporal gyrus (MTG) (15) where 75 transcriptomic distinct cell types were previously identified, including 45 inhibitory neuron types and 24 excitatory types as well as 6 non-neuronal cell types. A set of 142 marker genes are used to differentially distinguish the MTG cell types in (15). Structurally these genes form a highly differentially stable group (DS=0.734, p<8.66E-07), indicating strong cell type specificity, and 30 of these are among the disease genes, several uniquely associated with a disease (Suppl. Table 6).

We measure the tendency for disease gene co-expression to enrich in a specific cell type, using the Tau-score (τ) defined in (42) (**Methods**). For a gene *g,* 0 ≤ τ_g_ ≤ 1, measures the tendency for expression to range from uniform to concentrated in a specific cell type. Averaging τ over sets of genes representing a given disease, we obtain a measure of cell type specificity of each disease within MTG (**Suppl. Fig 14C**). While expression values between brain and non-brain disease genes differ only marginally (p=0.005), there is a highly significant difference in τ specificity between these groups (p < 2.2 *x* 10^-16^) (**Suppl. Fig 15A,B**) indicating specialized cell type involvement in genes associated with brain diseases (**Suppl. Fig. 15C**). Pooling to the 7 phenotypic categories (**Fig. 3B**), the classes psychiatric (2.52 *x* 10^-74^), movement (1.71 *x* 10 ^11^), and substance abuse disorders (3.58 *x* 10 ^11^) show the highest cell type specificity, while tumors, developmental disorders and neurodegenerative diseases to a lesser degree.

**Figure 3.**
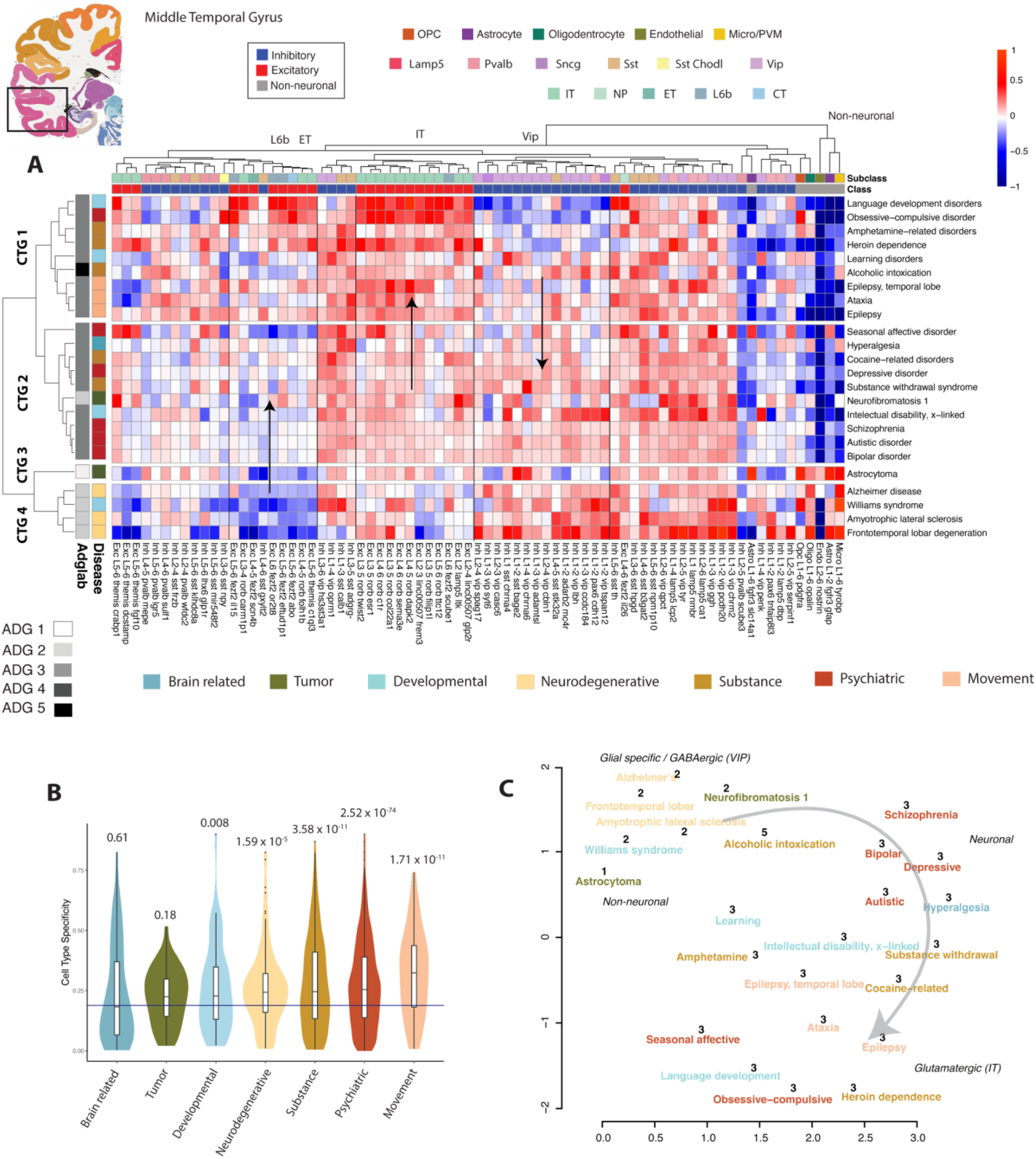
Disease genes and cell types of middle temporal gyrus. Reference plate from the *Allen Human Brain Atlas* (http://human.brain-map.org) containing middle temporal gyrus (MTG) region. A) Mean cell type expression (CPM) of 24 cortex related brain diseases (**Methods**) of 15,928 MTG nuclei over 75 cell types identified in (15). Diseases and cell types are clustered and identifies four cell type groups **CTG 1-4** based on cell type expression enrichment. Left annotation: **ADG** group membership determined by **Fig. 1**, and GBD phenotypic classification. Top annotation: Major cell type classes (excitatory, inhibitory, non-neuronal) and subclass level inhibitory (*Lamp5*, *Sncg*, *Vip*, *Sst Chodl*, *Sst*, *Pvalb*), excitatory (*L2/3 IT*, *L4 IT*, *5 IT*, *L6 IT*, *L6 IT Car3*, *L5 ET*, *L5/6 NP*, *L6 CT*, *L6b*), and non-neuronal (OPC, Astrocyte, Oligodendrocyte, Endothelial, Micro-glial/perivascular macrophages). Color coding is by class and subclass types. Arrows indicate increasing and decreasing cell type expression gradients. B) Cell type specificity τ measure pooled to phenotypic categories shows psychiatric and movement classes as most cell type specific. Bar: mean specificity over all cells, p-values of phenotype group shows significance. C) UMAP of combined mesoscale and cell type disease relationships color coded by phenotype (**Methods**). Numbers show original ADG membership with primary cell type annotation and excitatory gradient.

**Figure 3A** presents the clustering of mean expression profiles across the 24 cortical brain diseases. Diseases are now clustered by cell type specific expression and cell types and with annotations showing subclass level types (Inhibitory: *Lamp5*, *Pvalb*, *Sst*, *Sst Chodl*, *Vip*; Excitatory: *IT*, *NP*, *ET*, *CT*, *L6b*; and 5 non-neuronal types.) Cell type analysis in **Fig. 3** identifies four distinct *Cell Type Groups* (**CTG 1-4**) for these diseases. Here, **CTG 1**, representing several movement and substance abuse disorders, is characterized by a strong enrichment of neuronal excitatory IT over inhibitory Vip cell types (p < 5.53 *x* 10^-12^), and low expression of non-neuronal types.

**CTG 2**, representing several phenotypic classes and the major psychiatric (48) diseases, is higher in subclass *Vip* expression, exhibits more balanced pan-neuronal expression, and is also low in non-neuronal types. **CTG 3**, representing the non-neuronal enriched tumor-based diseases, is similar in profile to CTG 2 but with pronounced non-neuronal expression and captures ADG 1 diseases from the whole brain analysis. Finally, **CTG 4** has predominant enrichment in inhibitory neurons over excitatory (p < 1.343 *x* 10^-13^) and specialized non-neuronal types associated with the neurodegenerative diseases. Cellular level analysis consistently separates the stronger cortex expressing groups ADG 2-4 from the whole brain analysis (**Fig. 3A**, left annotation).

The basic cell types (inhibitory, excitatory, non-neuronal) of **Fig. 3** differentiate major disease groups of **Fig.1**, corroborating the module-based analysis of **Fig. 2C**. Expression clusters both at the subclass type level *Vip*, *Sst*, *Pvalb*, *IT*, *L6b*, and non-neuronal types **(Suppl. Fig. 16**), and analysis of variance at fixed cell types (**Suppl. Fig. 17**) shows that the highest variation across diseases occurs for excitatory and non-neuronal types. Further, **Fig. 3** illustrates gradients of increasing expression in excitatory cell types from **CTG 1-4** (CTG 3-4 p<0.0623, CTG 2-4, p < 3.56 *x* 10^-9^, CTG 1-4, p < 2.93 *x* 10^-18^) in *IT*, *ET* and *L6b* cell types across CTG with particular enrichment in language development, heroin dependence and obsessivecompulsive disorders (43). While inhibitory variation as a class is not significant across cell type groups, vasoactive intestinal peptide-expressing (*Vip*) interneurons show, by contrast, a decreasing gradient in expression from CTG 1-4 (CTG 1-2, p < 4.09 *x* 10^-10^,CTG 1-4 p < 8.26 *x* 10^-11^,CTG 2-4, 0.0006). Here pronounced enrichment of *Vip* interneurons, regulating feedback inhibition of pyramidal neurons (44), is seen in Alzheimer’s disease (45), frontotemporal lobar degeneration, ALS (46), and Williams syndrome (47). This gradient based analysis provides a means for distinguishing these cortically expressing diseases.

There is consistency between the structural (**Fig. 1**) and cell type analysis (**Fig. 3**) and their grouping by phenotypic class, despite data being limited to nuclei from a single cortical area (**Suppl. Figure 18**). We therefore combine the mesoscale and cell type approaches, averaging disease gene expression correlation matrices for 24 cortical diseases (**Methods**) and forming a consensus UMAP **Figure 3C** that graphically illustrates the transcriptomic landscape of major cortical expressing brain diseases, with key congruences and differences with phenotype association. The embedding in **Fig. 3C** shows grouping by original **ADG**, colored by phenotype, with labelling of primary cell types, and the significant excitatory cell type gradient in cortical expression.

The primary psychiatric diseases autism, bipolar disorder, and schizophrenia exhibit a largely similar expression profile **Figure 3A**, but detailed variation is overshadowed by stronger variation in other disease groups, and by the large number of genes associated with these three diseases. These disorders with a heritability of at least 0.8, are amongst the most heritable psychiatric disorders, and show a significant overlap in their risk gene pools (48). We formed a covariance matrix of cell type expression thresholding for significance(**Methods**). Interestingly, excitatory variation dramatically exceeds inhibitory and non-neuronal for these diseases (49) and accounts for 70.7% of significant cell type interactions (**Figure 4A** and inset). There are significant covarying cell types unique to autism, bipolar, and schizophrenia (*Aut*, *Bip*, *Scz*), as well as to specific to pairs of diseases (*Aut-Bip*, *Aut-Scz*, *Bip-Scz*). In particular, we found Aut-Scz (magenta) interactions with cell types of superficial layers (*Linc00507 Glp2r*, *Linc00507 Frem3*, *Rorb Carm1p1*), *Bip-Scz* in intermediate layer types (*Rorb Filip1*, *Rorb C1r*), and a unique enrichment of bipolar risk gene expression in *Rorb C1r*. Remarkably, although the genes enriched in a given cell type differ between the three disorders (**Suppl. Figure 19**), specific neuronal circuits are shared between the diseases (50, 51). **Figure 4C** derives associated biological processes and pathways of the unique *Aut*, *Bip*, *Scz* genes (g=19,20,25) that have significant expression in the interaction map of **Fig. 4B**. (**Suppl. Table 7**). The graph illustrates differential phenotype, with genes uniquely associated with autism linked to brain development, schizophrenia-associated enriched genes are implicated in dendritic outgrowth, and bipolar-associated genes are linked to circadian rhythm (52). The expression of these unique genes have distinct profiles across the implicated cell types, with schizophrenia exhibiting pan-excitatory expression (**Suppl. Fig. 18**). Cell type-specific interrogation of risk gene expression profiles provides insight into how polygenic risk impacts distinct types of neurons and neuronal circuits.

**Figure 4.**
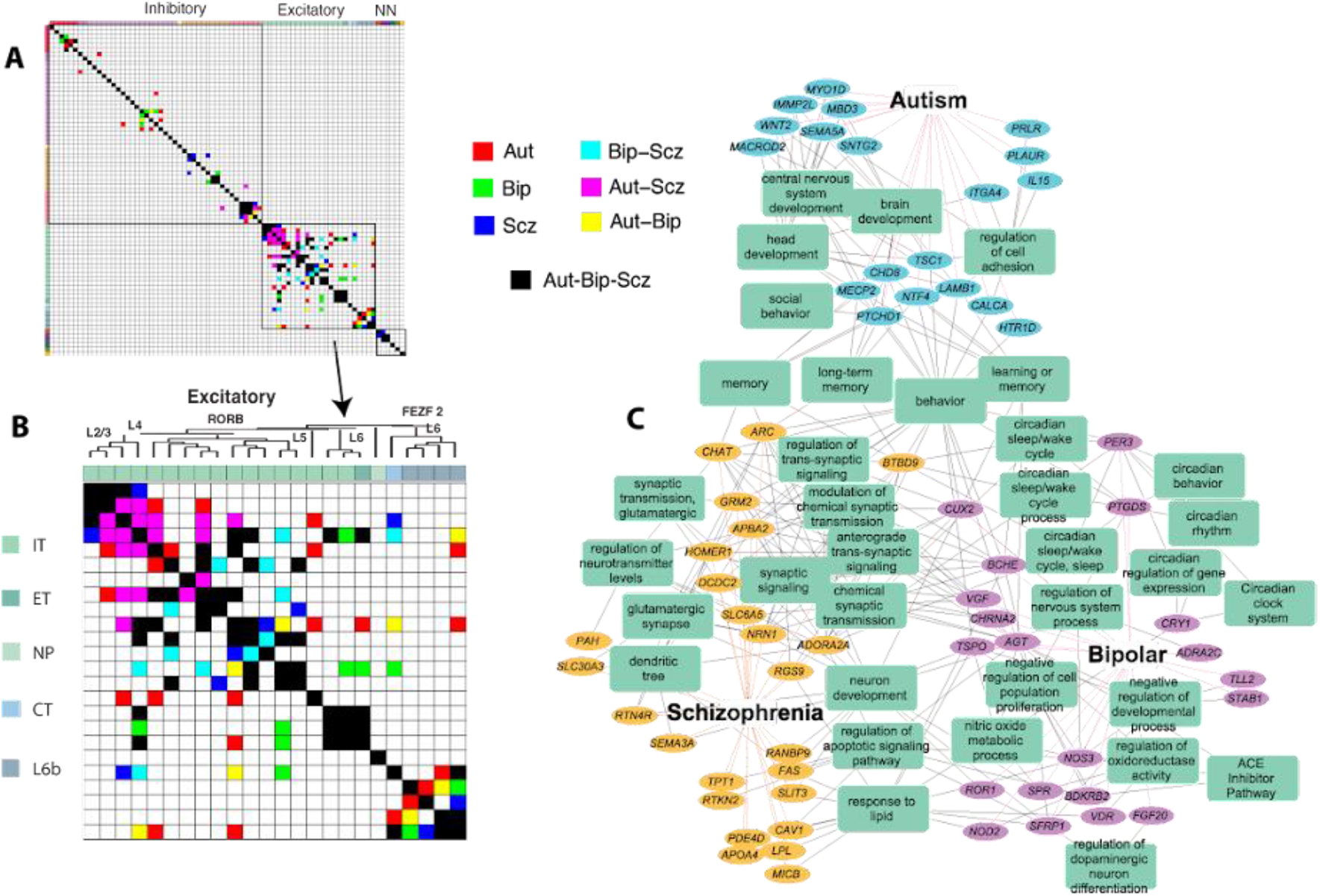
Cell type profile of autism, bipolar, and schizophrenia in human MTG. A) Significant cell type-specific covariation of gene expression for major psychiatric disorders (**Methods**). A) All 75 cell types with magnification B) of 24 excitatory types, color coded by disease combinations. Autism (*Aut*, red), bipolar disorder (*Bip*, green), and schizophrenia (*Scz*, blue) interactions unique to these diseases, *Aut-Bip* (cyan), *Aut-Scz* (magenta), and *Bip-Scz* (yellow) unique to pairs, *Aut-Bip-Scz* all. Excitatory cell types (*IT*, *ET*, *NP*, *CT*, *L6b*) and dendrogram from (15). C) Cell type-specific genes unique to *Aut, Bip, Scz* from (B) and representative enriched biological processes and pathways.

### Brain diseases in mouse and human cell types

Single cell profiling enables the alignment of cell type taxonomies between species, analogously to homology alignment of genomes between species. To examine conservation of disease-based cellular architecture between mouse and human, we used an alignment (15) of transcriptomic cell types from human MTG to two distinct mouse cortical areas: primary visual cortex (V1) and a premotor area, the anterior lateral motor cortex (ALM). This homologous cell type taxonomy is based on expression covariation and the alignment demonstrates a largely conserved cellular architecture between cortical areas and species identifying 20 interneuron, 12 excitatory, and 5 non-neuronal types (**Fig. 5A**.) We use this alignment to study species specific cell type distribution over the 24 cortex disease groups both at resolution of broad cell type class (N=7, e.g., excitatory), and subclasses (N= 20, as in **Fig. 3**) where non-neuronal cell types are common between both levels of analysis.

**Figure 5.**
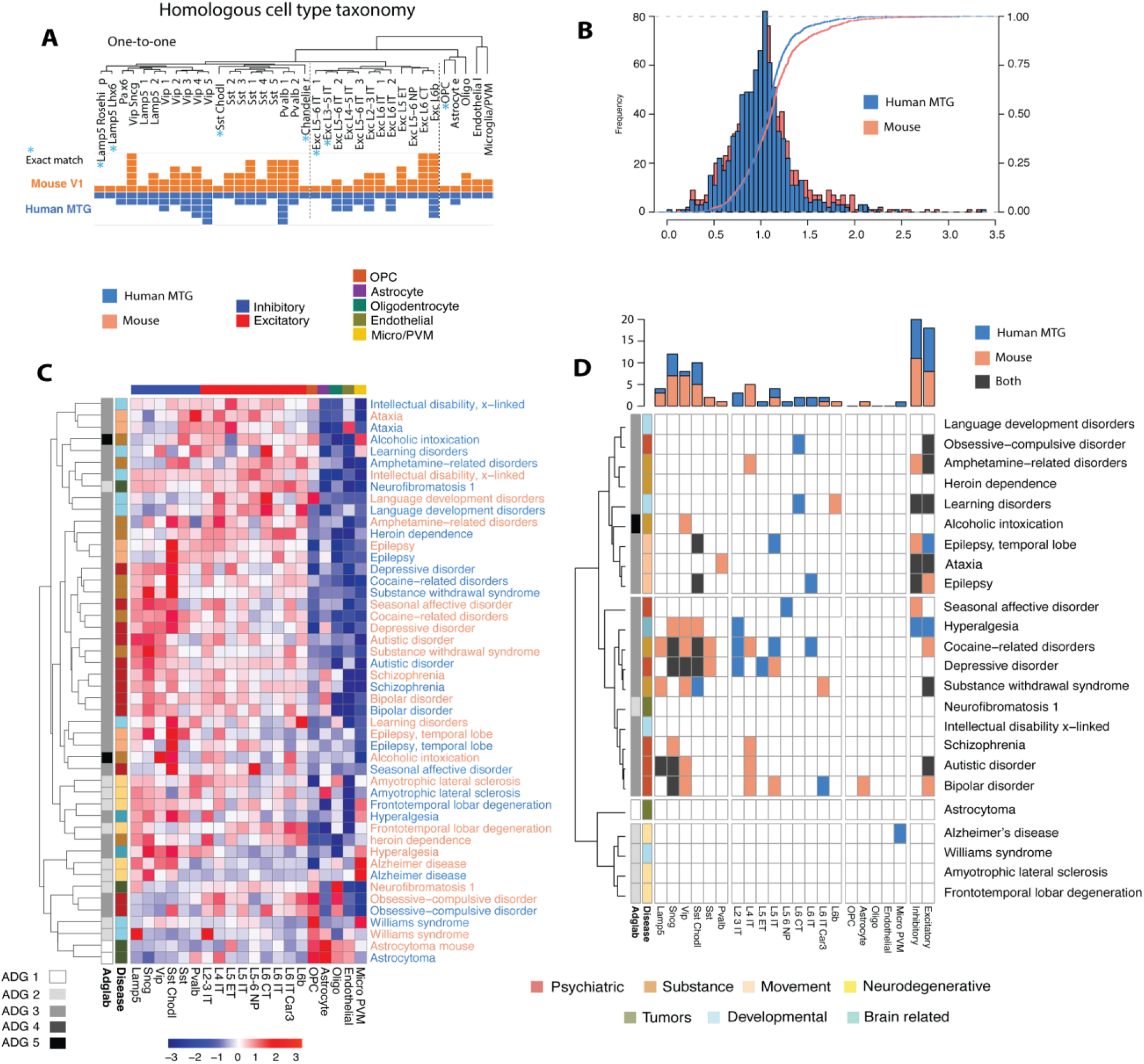
Disease based cell type expression in mouse and human. A) Alignment of transcriptomic cell types obtained in (15) of human MTG to two distinct mouse cortical areas, primary visual cortex (V1) and a premotor area, the anterior lateral motor cortex (ALM) B) Histogram of expression-weighted cell type enrichment (EWCE) values (42) human and mouse over subclass level of 20 aligned cell types. K-S goodness of fit test (**Methods**) shows that the distributions are marginally distinct (D=0.091, p=0.035). C) Simultaneous clustering of mouse and human EWCE disease signatures at subclass level 6 inhibitory, 9 excitatory, 5 non-neuronal (orange: mouse, blue: human) shows consistent signature across species. Annotation top major cell classes, side disease phenotype and ADG membership. D) Significant species distinct EWCE by disease and cell type with color code (blue: human, orange: mouse, black: both species). Disease clustering is as in **Fig. 3** with the same annotations. Top barplot: number of cell type enrichments by species.

To identify cell type differences in brain disorders between mouse and human cell types we used expression-weighted cell-type enrichment (EWCE) analysis (42). Briefly, EWCE compares expression levels of a set of genes associated with a given disease to the genomic background, excluding disease-related genes, by performing permutation analysis (**Methods**) and thereby defining the probability for the observed expression level of the given gene set compared with random sets of genes. In contrast to the *tau* measure (**Suppl. Fig. 14**) which examines each gene separately and measures only divergence of cell type expression from uniformity, EWCE evaluates all genes in a disease *simultaneously* and identifies concentration of cell-type expression for the group. The correlation of EWCE values aligned between mouse and human (**Suppl. Fig. 20**, ρ=0.633), is reflective of broadly conserved expression patterns (13) with no minimally significant difference in EWCE distribution (**Fig. 5B**). More remarkably, simultaneous clustering of EWCE mouse and human aligned cell types (**Figure 5C**, orange (mouse), blue(human)) shows highly conserved cell type signatures at the subclass level across species for many diseases (median ρ=0.645) and shows that original ADG groups are preserved in the mouse. **Fig. 5B** shows that the subclass expression signature for ataxia, epilepsy, bipolar disorder, ALS, Alzheimer’s disease, and schizophrenia, are more similar across species than to any other disease signature within species.

Cell type specific enrichment by EWCE corroborates specificity of major cell types and subclasses in both mouse and human. Psychiatric and substance abuse dominate the inhibitory (64%) and excitatory (70%) enrichments, consistent with the τ-specificity of **Fig. 3B**. **Fig. 5C** presents a more detailed view of the differences between mouse and human cell type enrichments. Here we find no significant enrichments in either species for several diseases including astrocytoma, neurofibromatosis 1, and frontotemporal lobar degeneration, while inhibitory subclasses *Lamp5*, *Sncg*, *Vip*, *Sst Chodl* show enrichment in both species (*Sst Chodl*, cocaine; *Sncg*, autistic, bipolar). Unique human enrichments are far more common in excitatory subclasses (*L6 IT Car3*, bipolar, *L2/3 IT*, *L5 ET*, depressive, *L6 CT*, learning disorders), and the only unique non-neuronal enrichment found is in human microglia/PVM for Alzheimer’s disease (p<0.0012). Although distribution of disease implicated cell types is largely conserved, **Fig. 5C** identifies several species-specific differences.

## Discussion

We presented a brain-wide molecular characterization of common brain diseases from the perspective of neuroanatomic structure, aiming to describe major transcriptomic relationships that vary with common phenotypic classification. Precise phenotypic classification of diseases is challenging due to variations in manifestation, severity of symptoms, and comorbidities (10, 53). We used the *Global Burden of Disease* (GBD) study from the *Institute for Health Metrics and Evaluation* (www.healthdata.org) for high-level phenotypic categorization, as this work is a continuously updated, globally used, comprehensive, and a data-driven resource. While described at coarser phenotypic resolution, such analysis is valuable for understanding the overall molecular relationships between common diseases and anatomic patterning of their gene expression in the brain, and to hint at interactions between genes for potential translational follow up.

For disease associated genes, *DisGeNet* is one of the largest resources integrating human disease genes and variants from curated repositories and provides a standard approach to select genes for the study. Determining implicated genes in disease states presents considerable uncertainty, and any study is likely to miss important associations. In particular, notably absent from our analysis are cerebrovascular diseases that account for the largest global burden of disability (10), and this limitation is due to relative under-sampling of rare vascular cell types in the *Allen Human Brain Atlas*. However, the approach presented is flexible and data driven and can be readily updated with other diseases of interest or associated genes following the steps in our accompanying *Jupyter* notebooks. As cell type data is now being generated in multiple regions of the human brain through the *Brain Initiative Cell Census Network* (BICCN, www.biccn.org) and upcoming *Brain Initiative Cell Atlas Network* (BICAN) this work can be readily extended.

Transcriptomic patterns of brain diseases cluster spatially into five major disease groups (**ADG 1-5**), largely recapitulated using cell type data from a single cortical area. **ADG 1** consists primarily of pan-glial diseases including most brain tumors, multiple sclerosis, migraine, and certain dementias and are transcriptomically distinct (**ADG 1**). Most neurodegenerative diseases (**ADG 2**) involve common neuronal (particularly cortex and hippocampus) and glial patterning (**Figs. 1A**, **2C**) effectively distinguishing them from largely glial based **ADG 1**. **ADG 3** shows the strongest neuronal patterning of all four disease groups with pronounced expression within the telencephalon, with minimal glial expression. This group mainly consists of psychiatric and substance abuse disorders, and epilepsies, recapitulating the known close genetic relationship between these disease groups (54). **ADG 4,5** comprise a combination of GBD diseases with modest cortical expression, enriched in glutamatergic cell types, and with a larger number (26%) of anatomic structural markers in basal ganglia, hypothalamus, and lower brain structures, **ADG 5** is distinguished from **ADG 4** by strong expression in the thalamus. The general association of these disease groups is reproduced both in cell type specific analysis of middle temporal gyrus and corroborated in the mouse.

This study finds diverse phenotypes and clinical presentations have shared anatomic expression patterns and may provide insight into disease mechanisms and frequency of comorbidity. For example, language development disorders, OCD and epilepsy, temporal lobe are phenotypically diverse, yet all belong to **ADG 3**, and cell type analysis in **Fig. 3A** illustrates a correlated cell type signature with strong IT excitatory subclass expression. While these are broad categorizations, there is reproducible structure to anatomic disease profiles illustrated through differential expression stability analysis and through correspondence between mouse and human cell type profiles (**Fig. 5C**). The use of brain wide relationships to study and characterize brain disorders complements localized expression of disease genes and their coregulation that will be potentially invaluable, especially as large-scale spatially resolved cell type studies become available.

The general correspondence of structural and cell type approaches even when restricted to a single cortical area (MTG) suggests a consensus organization and amplifies the value of cell type and tissue-based deconvolution methods, particularly when extrapolating these results to multiple brain regions. An intriguing finding is how diseases associated with pronounced cortical expression are organized along a gradient of excitatory cell types. This organization, also anti-correlated with an inhibitory gradient of specialized subclass interneurons, potentially provides insight into new methods for classifying brain diseases. Cortical spatial gradients of gene expression were first observed in earlier tissue-based studies (14) and although originally attributed to sampling resolution, have been now observed at cellular resolution (20, 55). With increasing scale of single cell studies this may provide an important means of disease comparison that clarifies phenotypic associations.

This work is complementary to studies on shared genetic heritability of common disorders of the brain. The *Brainstorm* consortium studied a large cohort GWAS meta-analysis demonstrating that common genetic variation contributes to the heritability of brain disorders, and showing that psychiatric disorders share common variant risk, with other neurological disorders appearing more distinct from one another and psychiatric disorders (56). This result is also seen in the present study, with a lower transcriptional variance in both structural and cell type profiling between schizophrenia, bipolar and autistic disorders compared with a wider range of anatomic patterning in neurological disorders (**ADG 4,5**). A striking finding however is the variability of excitatory cell types in psychiatric diseases, and certain species-specific expression differences in these psychiatric and substance abuse (**Fig. 5B**). While there have been several lines of evidence that inhibitory cell types are impaired in psychiatric disorders (57, 58) (e.g., depression, bipolar disorder, and schizophrenia), results here indicate that excitatory pathways may be equally important. There are, however, limitations to a cell type enrichment approach. Some diseases may involve gene pathways shared across cells rather than involvement of subsets of cell types or brain regions, and as others have found, cell type enrichment of disease genes here does not necessarily match cell types with expression differences in disease vs. control tissue (59, 60). Exploring the transcriptomic architecture of these disorders is a fully new field that has been underexplored and these findings support the transcriptomic hypothesis of vulnerability that in polygenic disorders, genes that are co-expressed in a certain brain region or cell type are much more likely to interact with each other than those that do not follow such a pattern (11, 12).

While previous work has shown conservation of neuronal enriched expression patterning between the mouse and human (13, 16), a recent novel alignment of mouse and human cell types in middle temporal gyrus now allowed for a more specific analysis. For example, microglial involvement in Alzheimer’s disease is well established, seen in **Fig. 3**, and found uniquely human enriched (**Fig. 5B**). Here we show that the mouse appears to be evolutionarily sufficiently close to identify potentially relevant cell types and a striking conserved signature across subclass cell types for many diseases. This is important as it suggests we can leverage cross species cell type atlases to indicate disease risk gene patterning (61). While homology alignment of cell types between mouse and human may provide insight into convergent mechanisms based on species-specific differences, further human data is needed to implicate disease genes with cell function. We are aware of the different sampling protocols and numerous complications of cross-species comparison and our work addresses broad trends across diseases and cell types as opposed to differential expression of specific genes. Our results describe the structural and cellular transcriptomic landscape of common brain diseases in the adult brain providing an approach to characterizing the cellular basis of disorders as brain-wide cell type studies become available.

## Methods

### Disease genes

To obtain the gene disease associations, we used the DisGeNET database (21), a discovery platform with aggregated information from multiple sources including curated repositories, GWAS catalogs, animal models and the scientific literature. DisGeNET provides one of the largest gene-disease association collections. The data were obtained from the April 2019 update, the latest update related to the gene-disease association at the time of analysis. An original list of 500 diseases with connection to the brain were intersected with the provided repository at DisGeNET. For each disease, the main variant was selected, and rare familial/genetic forms were not included in the analysis. For this study, we included genes with gene-disease association reported at least in one confirmed curated (i.e., UNIPROT, CTD^TM^, ORPHANET, CLINGEN, GENOMICS ENGLAND, CGI, PSYGENET) (for details see https://www.disgenet.org/dbinfo). Since the goal of the study is to investigate the similarities and distinctions between brain-related disorders, disorders with less than 10 genes associated with them forforwere excludedrom the analysis. Finally, 15 disorders of peripheral nervous system or a second level association to the brain (e.g., retinal degeneration) were removed. This procedure resulted in 40 brain disorders with their corresponding associated genes. Finally, for these 40 disorders, we performed a literature review of the current GWAS studies to add all the missing genes from the DisGeNET dataset. These 40 diseases include brain tumors, substance related, neurodevelopmental, neurodegenerative, movement, and psychiatric disorders (**Supp. Figure 1**).

### Dataset description

Anatomic based gene expression data was extracted from 6 post-mortem brains (14). The extracted samples were divided into 132 regions based on the anatomical/histological extraction regions. These 132 regions were further pooled/aggregated into 104 regions including cortex (CTX,8), hippocampus (HIP,7), amygdala (AMG,6), basal ganglia (BG,12), epithalamus (ET,3), thalamus (TH,10), ventral thalamus (VT,2), hypothalamus (HY,16), mesencephalon (MES, 11), cerebellum (CB,4), pons (P,8), pontine nuclei (PN,2), myelencephalon (MY, 12), ventricles (V,1), white matter (WM,2) (**Suppl. Table 3**). The resulting gene by region matrix was further averaged between subjects to produce one representative gene expression by region matrix. Each gene expression profile was further normalized across the brain regions. Cell type data in human is based on snRNA-seq from middle temporal gyrus (MTG) largely from postmortem brains (15). Nuclei were collected from eight donor brains representing 15,928 nuclei passing quality control, including those from 10,708 excitatory neurons, 4,297 inhibitory neurons and 923 non-neuronal cells. Cell type data from the mouse represents 23, 822 single cells isolated from two cortical areas (VISp, ALM) from the C57GL/6J mouse (18).

### Cell-type specificity

Calculated based on the Tau-score defined in (42). This measure has previously been employed using the same dataset (15). Briefly, Cell-type specificity τisτis defined as:

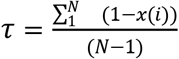

where, x(i) is the gene expression level in each cell-type for a given gene normalized by the maximum cell-type expression of that gene, and N is the number of the cell-types in the analysis.

### Disease-Disease similarity index

In order to calculate the similarity between each pair of disorders we used the gene expression patterns across 104 brain structures, removing overlapping genes from each pair of disorders during clustering. In presenting heatmaps the full set of genes in each disease are averaged. Distance metric between diseases is 1 - 〉 where 〉 is Pearson correlation between structure or cell type profile also removing common genes. The procedure for disease similarity using cell-type data used the gene expression pattern across the 75 cell-types (instead of brain regions) in human cells extracted from MTG. For clustering in both cases, we used agglomerative hierarchical clustering with Ward linkage algorithm (i.e., Ward.2 in R hclust function, R version 3.6.3)

### Gene Expression Differential stability (DS)

Gene expression differential stability was calculated for each gene as the similarity of its expression pattern across 6 post-mortem brains. For each pair of brains, the correlation of expression patterns across overlapping brain structures was calculated. The mean correlation over these 15 pairs was used as the differential stability for the given gene. (For more details see (13).

### Disease-module association

For each gene the relationship between gene expression for each gene and module is calculated as explained in (13)(i.e., each gene expression pattern is correlated with eigengene pattern from modules within each of 6 postmortem brains). These correlation values are then normalized using Fisher r-to-z transform and averaged across brains. For each module, the gene associations were then standardized (mean=0, 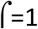). Finally, these values are averaged across genes associated with each disease to calculate the disease module association.

### Disease related gene expression within cell-types

We used expression-weighted cell-type enrichment (EWCE) analysis (https://bioconductor.riken.jp/packages/3.4/bioc/html/EWCE.html; (42) to identify cell types showing enriched gene expression associated with each of the 6 brain donors. Briefly, EWCE compares the expression levels of the genes associated with a given disease to the background gene expression (all genes, excluding the disease-related genes) by performing permutation analysis and defining the probability for the observed expression level of the given gene set compared against a random set of genes. We used N=100,000 as the permutation parameter and performed the analysis at two cell-type category levels. The two levels included broad cell-types (N=7) and cell-subclasses (N= 20). The non-neuronal cell types were common between the two levels of analysis. These two levels were selected due to the availability of the homologous cell types in mouse and human cell dataset. Finally for each disease, we used false discovery rate (*FDR*) correction for multiple comparisons for disease-cell type associations.

### Cell Type Specific Interaction and Functional Enrichment

Gene expression covariation across cell types is computed by absolute value of cosine distance similarity and then thresholded to 1.5σ. Functional enrichment analysis to identify significantly enriched (p-value <0.05 FDR Benjamini and Hochberg) ontological terms and pathways for unique disease gene sets was done using the ToppFun application of the ToppGene Suite (62). Representative enriched terms and genes were used to generate network visualization using Cytoscape application (63).

### Consensus Representation

Consensus UMAP was constructed by averaging pairwise gene set correlation matrices for structural and cell type data, and forming a 2D UMAP using R.

### Statistical Analysis

All statistical analysis and visualization were conducted in R (www.r-project.org), a Jupyter notebook reproduces all analysis. To examine the differences in mean expression level between ADG groups we performed ANOVA tests. This was followed by direct comparisons between ADG pairs using unpaired t-test. All results were corrected for multiple comparisons using Benjamini-Hochberg correction controlling the false discovery rate. To examine the stability of the gene expression profiles, we repeated our analysis across 6 brains and searched for the matching pattern in other subjects for any given brain across ADG and GBD disease groups. The gene expression DS within each GBD group was compared to the general DS of all other genes in the dataset using independent t-tests.

## Data Availability

All data used in this manuscript are publicly available. The gene disease association data can be downloaded from https://www.disgenet.org/. The large-scale anatomic transcriptional patterns can be downloaded from http://human.brain-map.org/ and cell type data is available at http://celltypes.brain-map.org/.

## Code Availability

The script alongside a notebook file and all necessary data files for producing the figures are provided at https://github.com/yasharz/human-brain-disease-transcriptomics.

## Acknowledgements

The authors thank Christof Koch, Liane Ong, Jay Schulkin, Stephen J. Smith, and Theo Vos for insightful and helpful discussions.

