## Supplementary Figures for "Structural and cellular transcriptome foundations of human brain disease"

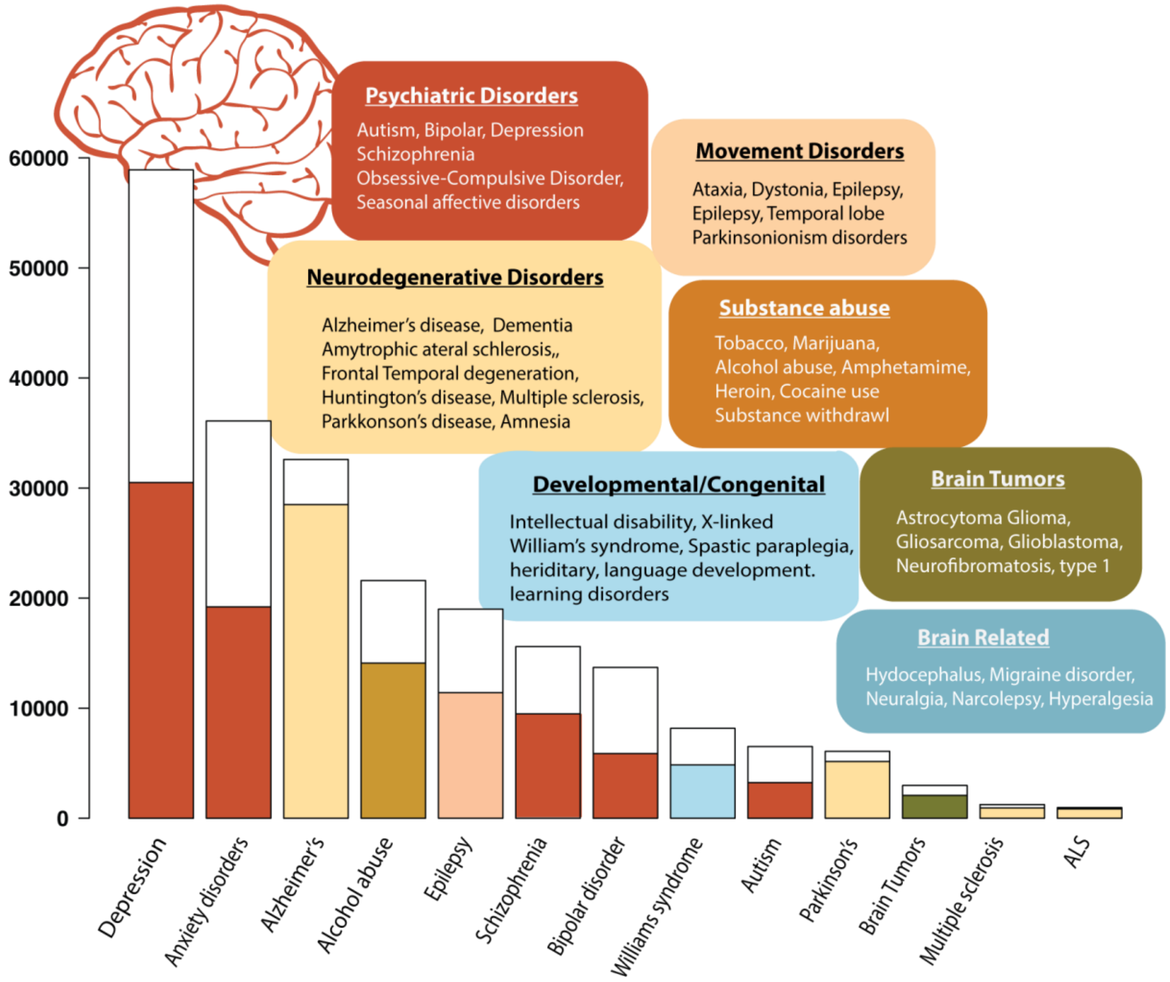


**Supplementary Figure 1. Classification and global burden of brain related diseases**. Major human brain diseases and classification according to the *Global Burden of Disease (GBD)* study (1, 2) partitioned by seven broad classes. The GBD study established the standard *Disability Adjusted Life Years* (DALY) metric to quantify disease burden defined as the years lost due to premature death plus years lived with disability. DALY scores are shown according to the 2019 study for several larger classes with error bars in white indicating minimum and maximum projected loss of life and healthy years. While cerebrovascular diseases including brain ischemia and infarction and related disorders dominate (global 2017 DALY 55.1 million, not shown), the combined toll of psychiatric disorders has nearly twice DALY (110 million). Neurodegenerative diseases account for less (38.2 million) primarily through older populations with Alzheimer's disease and related dementia (30.5 million) DALY. Color palette for these major GBD classes is used throughout the analysis.

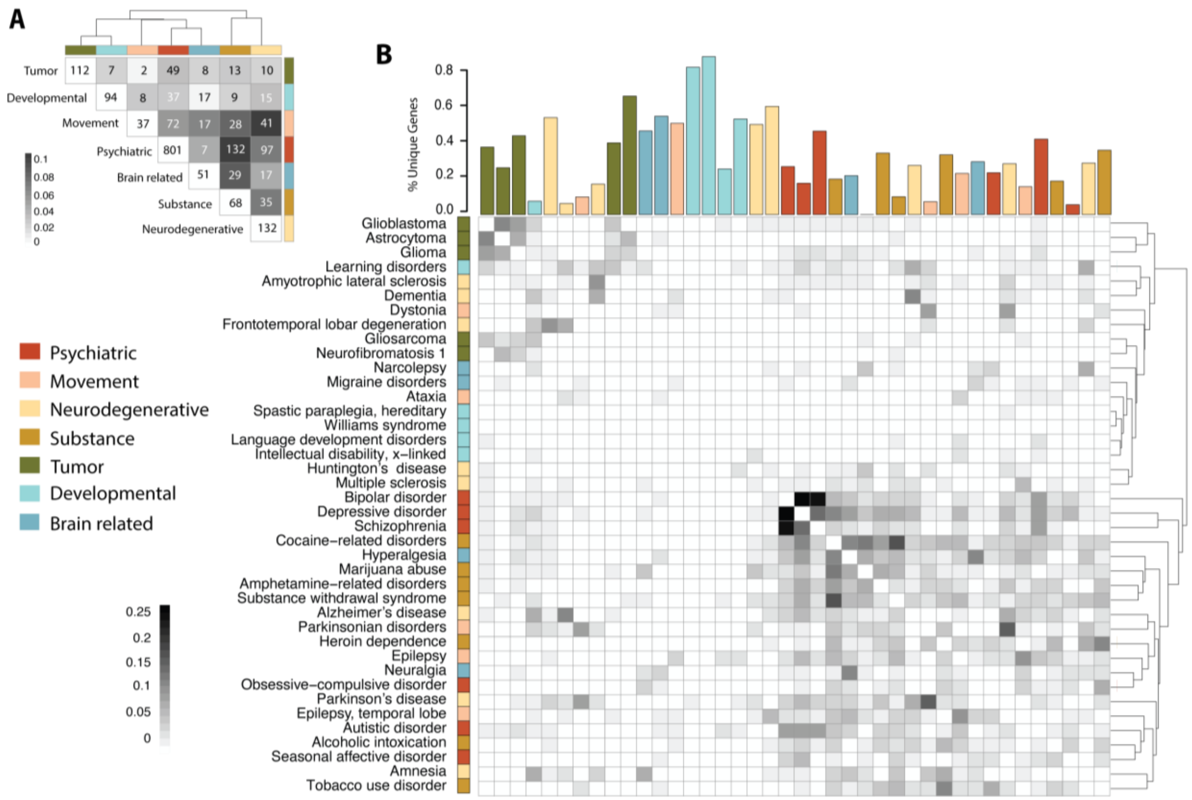


**Supplementary Figure 2. Neurological disorders and associated genes**. A) Jaccard clustering based on relative percentage of shared genes between GBD classes for disease genes in this study. Inset numbers: number of genes in intersection, with diagonal unique number to class. B) Similar clustering of 40 neurological diseases and disorders. Top panel: Fraction of genes uniquely associated with each disease. Color panel: membership GBD class for disease. Details of disease, gene sets, and metadata are given in **Suppl. Table 1**. Whereas the number of unique genes associated to GBD class psychiatric diseases (801) is 6 times larger than neurodegenerative diseases (132), a finer resolution does not reflect this bias with 110 genes (28.6%) unique to bipolar disorder whereas 31 genes (30.3%) are unique to Parkinson's disease, 59 (88.0%) unique to hereditary spastic paraplegia.

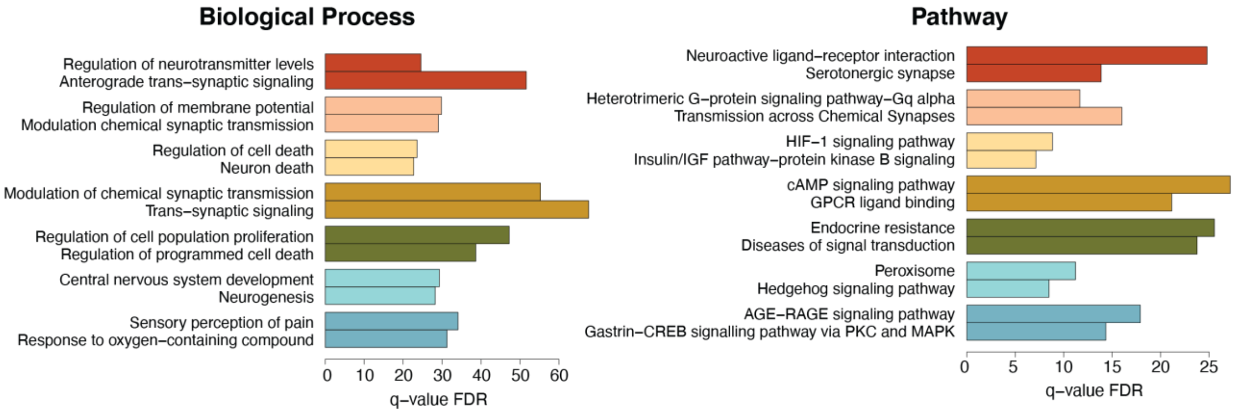


**Supplementary Figure 3.** **Biological process and pathway ontology analysis** (www.toppgene.org) of genes *uniquely* associated with major GBD classes reflect common identifying annotations for these disease classes measured by FDR q-value. Color code in legend for GBD classes is used throughout the analysis. Specific associations of interest include well-known alterations in synapse structure and function (FDR $q = 9.56 \times 10^{-50}$) (3), and abnormal levels of extracellular neurotransmitter concentrations (4) in several psychiatric and neurologic disorders ($q = 1.25 \times 10^{-22}$). Major depressive disorder is one of the most important mental disorders associated with altered serotonergic activity (5), with less clear association in schizophrenia (6) and addiction (7). Recent studies show that chronic type II diabetes mellitus (DM) is closely associated with neurodegeneration ($q = 2.07 \times 10^{-5}$), especially AD (8). The primary signaling pathway activated in insulin signaling is the phosphoinositide 3-kinase (PI3K)-protein kinase B (Akt) signaling stream, and defective IGF binding or IRS-1 signaling, as a result of insulin resistance, leads to cognitive decline in patients (9). Hedgehog (Hh) is one of few signaling pathways that is frequently used during development for intercellular communication, important for organogenesis of almost all organs in mammals, as well as in regeneration and homeostasis (10). This includes the brain and spinal cord and mutations in the human *SHH* gene and genes that encode its downstream intracellular signaling pathway cause several clinical disorders, include holoprosencephaly (11). Brain tumors and other cancers are strongly associated with defects in signal-transduction proteins (12), and cancers caused by certain viruses have contributed greatly to our understanding of signal-transduction proteins and pathways (13). Chronic morphine-induced molecular adaptation of the cAMP cascade has been confirmed in many and has been widely related to opioid dependence and withdrawal (14). These unique GBD class ontology annotations represent molecular function and pathways central to these major classes.

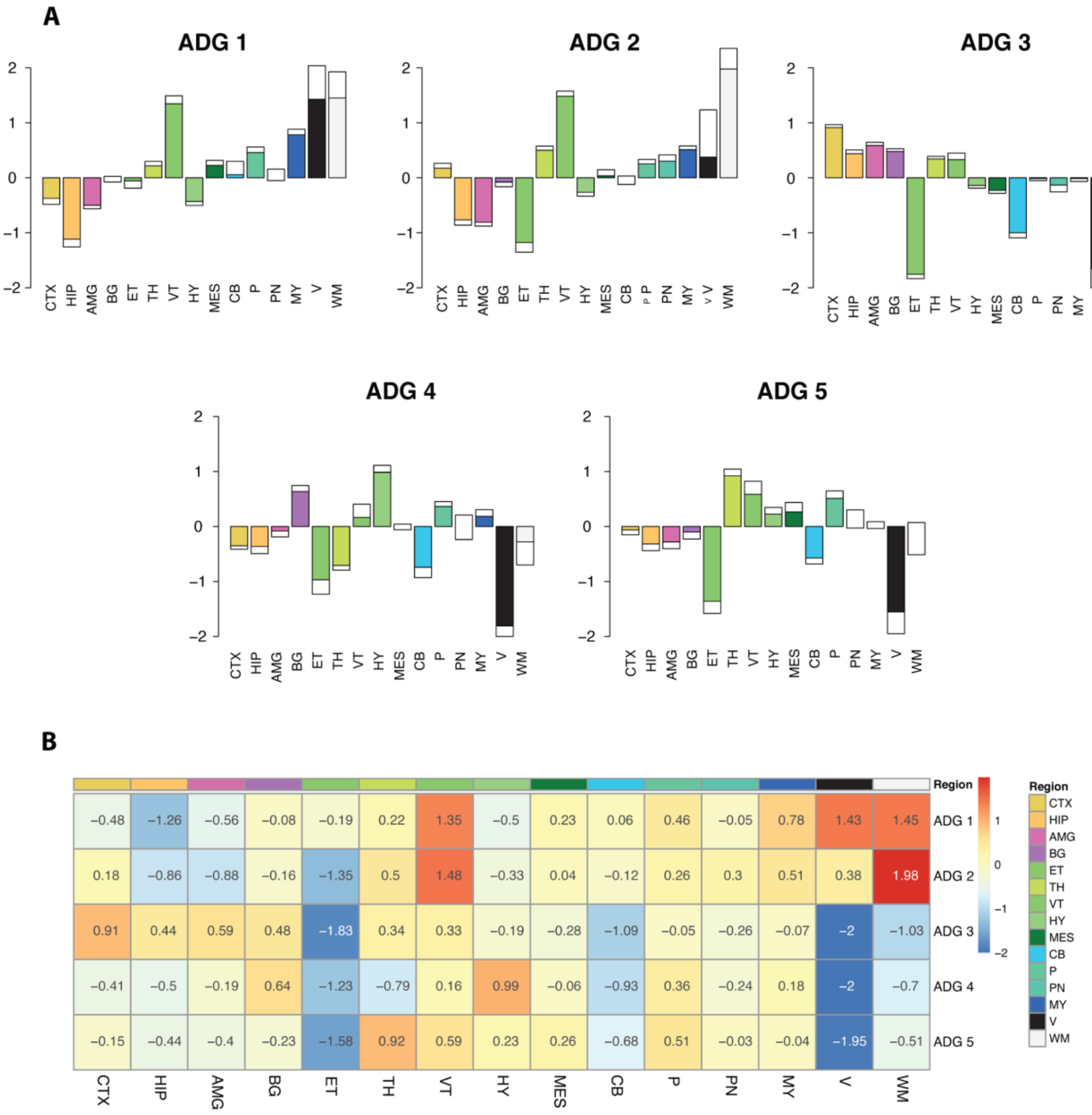


**Supplementary Figure 4. Mean anatomic transcriptomic profile for ADG groups.** A) Mean expression level of 15 major brain regions with standard error for each Anatomic Disease Group **ADG 1** – **ADG 5**. Major regions include cortex (CTX), hippocampal region (HPP), amygdala (AMG), basal ganglia (BG), epithalamus (ET), thalamus (TH), ventral thalamus (VT), hypothalamus (HY), mesencephalon (MES), cerebellum (CB), pons (P), pontine nuclei (PN), myelencephalon (MY), ventricles (V), white matter (WM). Color codes for regions are those used in Allen Human Brain Atlas (**Supplementary Table 3**) B) Mean z-scored expression level of 15 major structures summarizing major differentiating relationships across ADG groups.

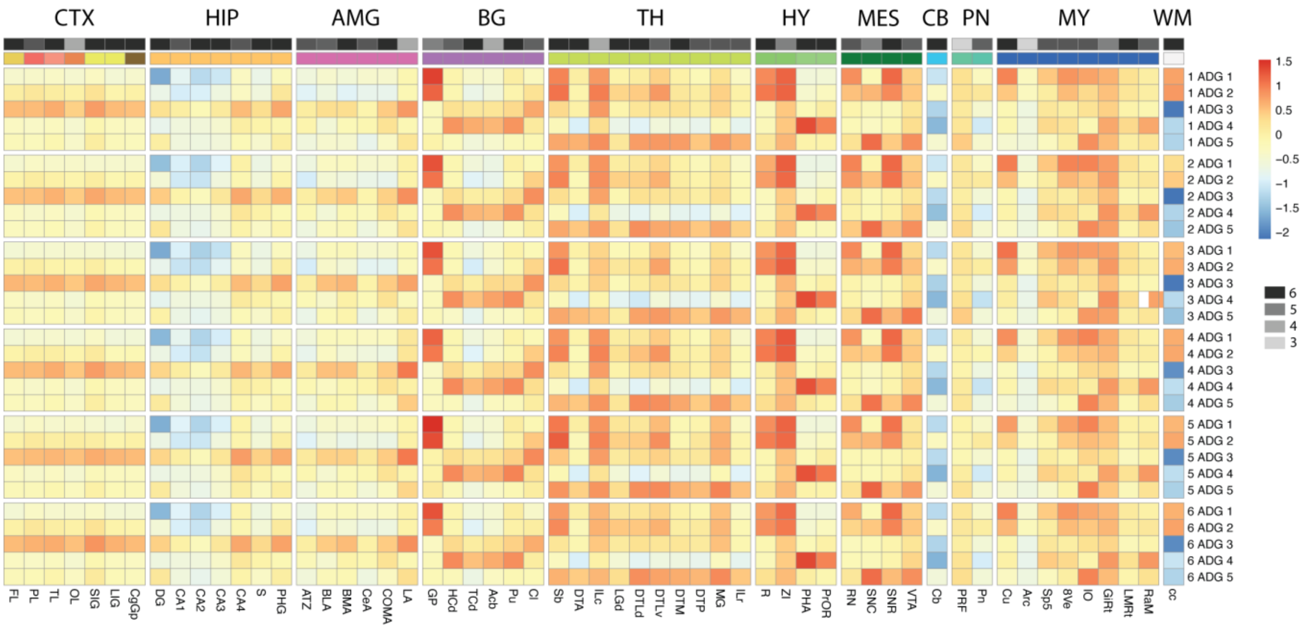


**Supplementary Figure 5. Reproducibility of ADG clustering.** A hold out analysis was conducted averaging the z-score normalized expression within each of the identified ADG groups identified in the full analysis of **Figure 1** with one of 6 brains data left out. On left 1 ADG 1 indicates that brain 1 data was removed and diseases in ADG groups averaged in the remaining 5 brains. Data is presented over 57 structures common to all 6 brains. Viewed as rows across structures, the reproducibility of expression patterning is seen to be highly consistent across hold out data sets with average correlation (ADG 1, ADG 2, ADG 3, ADG 4, ADG 5) = (0.983, 0.971, 0.976, 0.988, 0.977). Viewed as columns across structures the patterning has consistent differential expression across ADG groups. The annotation bar on top of the heatmap shows the maximum repeatable differential signature observed in each structure. The signature is exact (6) in all hold out brains for 27 structures and agree in all but one for 19 additional structures, only LA, PRF, and Arc displaying variability. The expression signature itself is computed and compared as follows. For each structure and each hold out dataset the z-scored expression values are rank ordered giving a permutation of 1,2,3,4,5 from lowest to highest across the **ADG 1-5**. Each expression pattern is assigned a unique integer **n** through unique prime factorization as **n**=$2^{(1)}3^{(2)}5^{(3)}7^{(4)}11^{(5)}$and these integers are tabulated to find the most occurring pattern across hold out brains. The maximum occurring signature 3-6 is shown in the annotation bar indicating similar conservation of signature to the hold out analysis, with 6 representing the exact relationship of ADG groups in all brains.

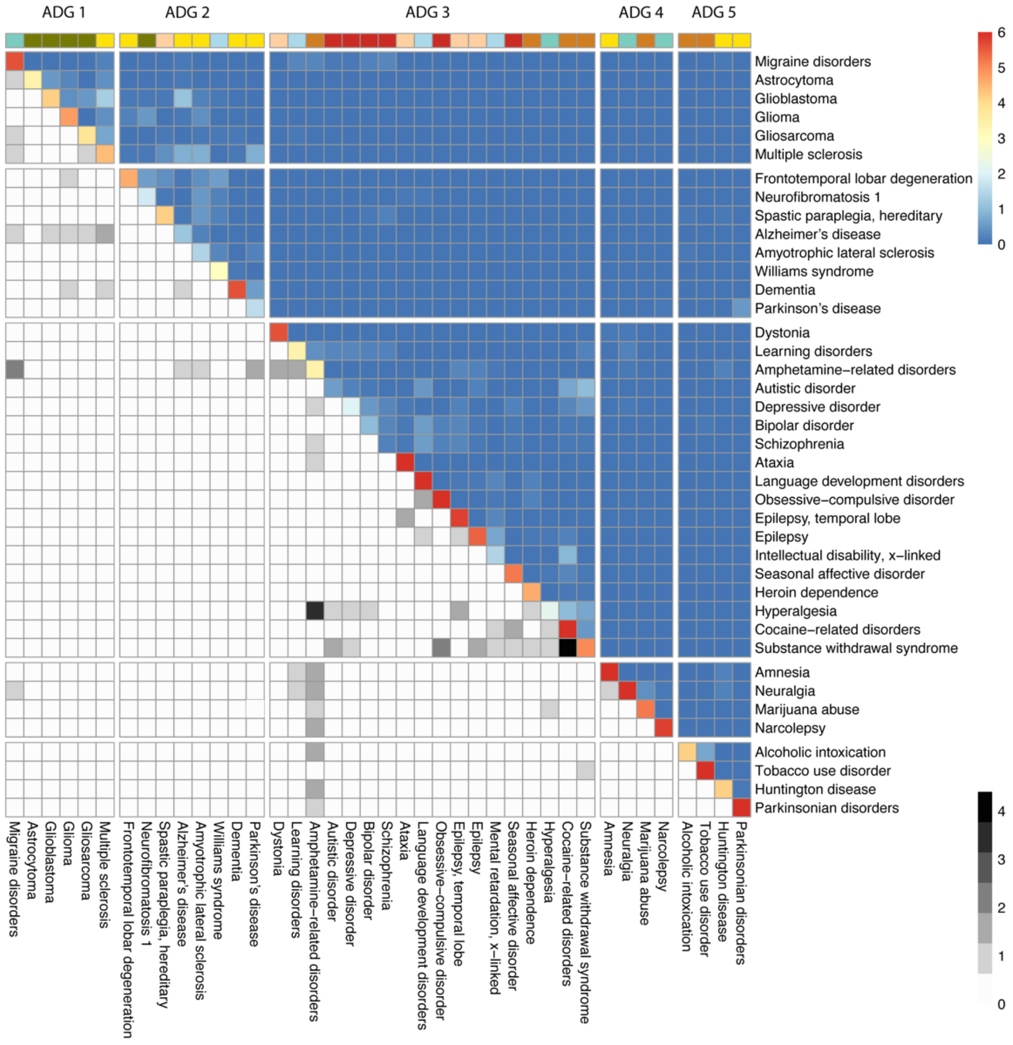


**Supplementary Figure 6. Holdout analysis and ADG.** (Diagonal and upper) In each of six Allen Human Brain Atlas (AHBA) subjects the mean disease transcription profile for each of 40 diseases across structures is computed and the most similar (Euclidean distance) disease in the remaining 5 subjects is identified. The upper diagonal matrix shows the distribution of identified diseases with key 0-6 indicating the number assignments to given disease. Thus, ataxia with score 6 has a transcriptomic profile more similar to ataxia for each brain than to any other disease in the remaining brains. Since the closest neighbor is an asymmetric definition, the average of the matrix and its transpose is presented. A majority 29/40 diseases are uniquely identified by majority voting. ADG groups 3, 4 and 5 have high identifiability across subjects while there is higher misclassification between ADG 1 and 2. Percent exact as in Figure 1C is ADG 1-5 (0.716, 0.537 0.644, 0.958, 0.875). Color bar shows Global Burden of Disease (GBD) groups. (Lower diagonal) A more stringent hold out analysis is conducted first eliminating common genes between the diseases as in Figure 1 and by seeking the closest disease in transcriptome profile other than the given disease. Here the distribution of disease mapping between brains is more variable having within ADG mapping ADG 1-5 (0.361, 0.187, 0.970, 0.175, 0.008)

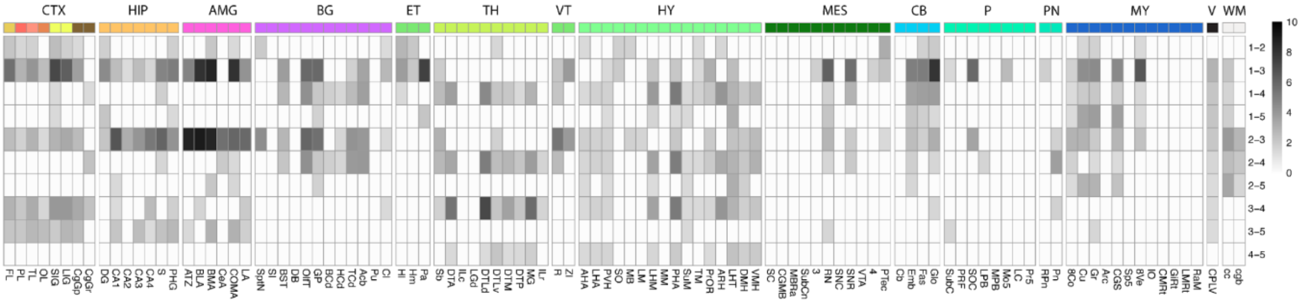


**Supplementary Figure 7. Pairwise comparison of ADG.** Pairwise B&H corrected (BH < 0.05) t-tests between **ADG** groups 1-5. Individual t-tests highlight the distinction in cortex expression between ADG 3 and other groups. The most significant structural ADG differences occur between **ADG 1-3** in cortex (frontal lobe (FL, $p < 2.71 \times 10^{-7}$)), short insular gyri (SIG $6.2 \times 10^{-9}$) , long insular gyri (LIG, $5.57 \times 10^{-8}$), in amygdala, basolateral nucleus (BLA, $1.8 \times 10^{-9}$), basomedial nucleus (BMA, $4.49 \times 10^{-10}$), in cerebellar nuclei, globose nucleus (Glo, $1.18 \times 10^{-9}$), and myelencephalon, vestibular nuclei (8Ve, $2.34 \times 10^{-8}$). **ADG 2-3** are distinguished in hippocampus, (CA1, $2.18 \times 10^{-8}$), subiculum (S, $8.31 \times 10^{-8}$), in amygdala (AMG), amygdalo-hippocampal transition zone (ATZ $1.94 \times 10^{-10}$ , BLA, $1.00 \times 10^{-10}$, BMA, $5.63 \times 10^{-10}$ ), and between **ADG 3-4** thalamus, anterior group of nuclei (DTA, $3.01 \times 10^{-7}$), lateral group of nuclei, dorsal division, (DTLv, $6.47 \times 10^{-9}$), and hypothalamus, posterior hypothalamic area (PHA, $1.21 \times 10^{-6}$).

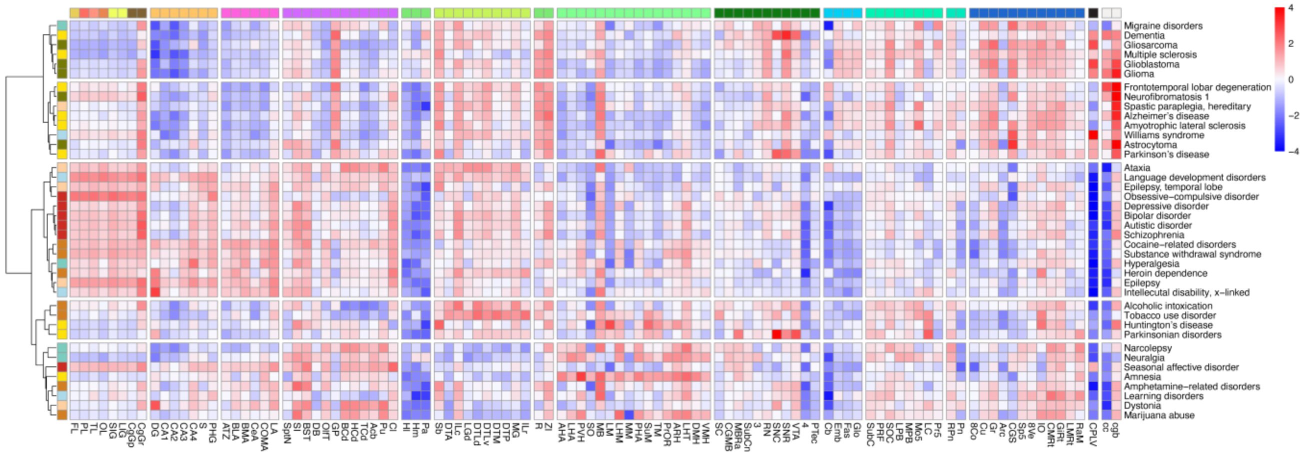


**Supplementary Figure 8.** Transcriptome patterning of 40 brain diseases with clustering *removing* pairwise overlapping genes also identifies five anatomic groups. Most distinctive is the strong match of **ADG 1** and **ADG 2** indicating the meaningful identity and distinction of these groups. Removing common genes retains the association of the majority of **ADG 3** psychiatric, substance abuse, and movement diseases. The grouping of diseases in **ADG 5** is identically preserved in the clustering, overall indicating common structure with **Figure 1.**

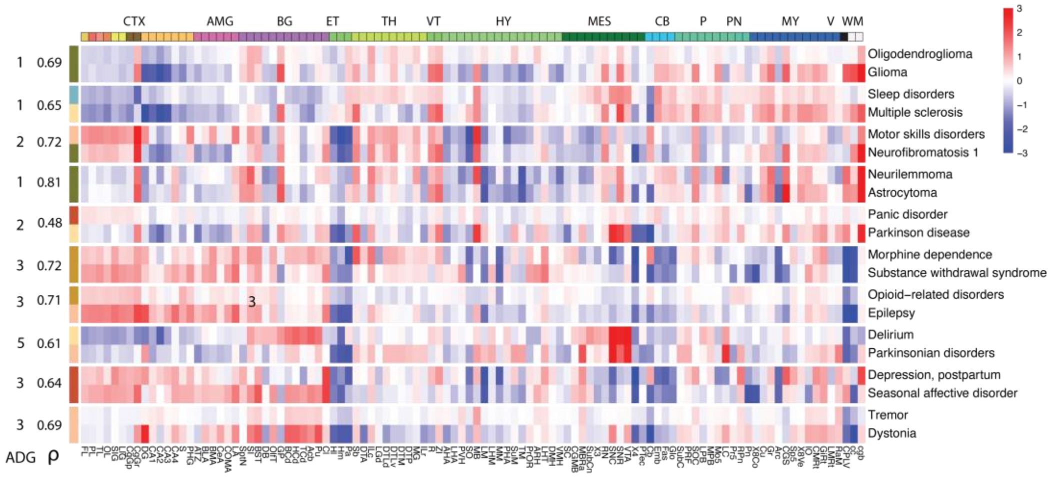


**Supplementary Figure 9. Anatomic transcriptome profile for limited gene diseases.** Ten diseases with limited associated gene sets (5 ≤ n < 10) and their relationship to ADG groups: Oligodendroglioma, sleep disorders, motor skills disorders, neurilemmoma, panic disorder, morphine dependence, opioid-related disorders, delirium, post-partum depression, and tremor. Right panel shows each disease and the closest correlated disease from the major set of 40 disorders. Left panel shows the RGB groups, the correlation with the closest disease from the major group, and its ADG group number. These small gene set associated diseases reflect the major ADG patterning observed in **Figure 1.** The profiles of oligodendroglioma and motor skills disorders further highlight the distinction between ADG 1 and ADG 2.

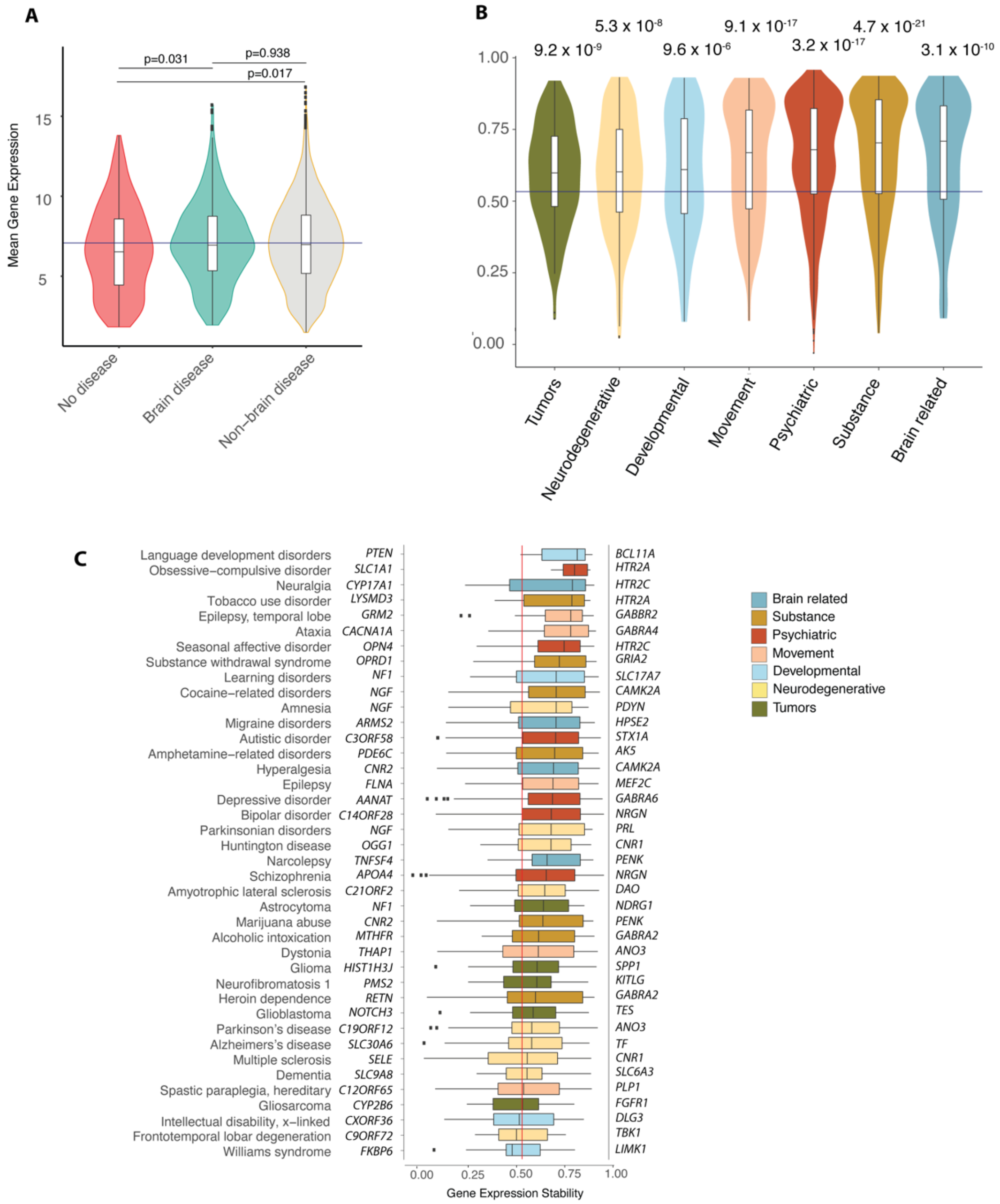


**Supplementary Figure 10. Expression levels of brain and non-brain diseases.** A) Expression levels of genes from Allen Human Brain Atlas (AHBA) classified as brain disease associated from this study (green), non-brain brain disease associated from OMIM study of (15) (gray) and remaining genes of AHBA not in these sets (red). Brain disease genes do not have significant expression differences from non-brain related genes, but both are different from non-disease associated genes with marginal significance. B) Distribution of DS by major Global Burden of Disease classes. Horizontal mean ρ=0.521 of 17,348 genes, with p-values shows significance (corrected for class size) of GBD mean differing from global mean. C) Disease gene stability for 40 diseases sorted by median DS; colors are GBD classification. Minimum and maximum stable genes for each disease are shown. DS: differential stability. The set of high DS genes annotated on right is substantially enriched for Gene Ontology biological processes and pathways compared to lower DS on left.

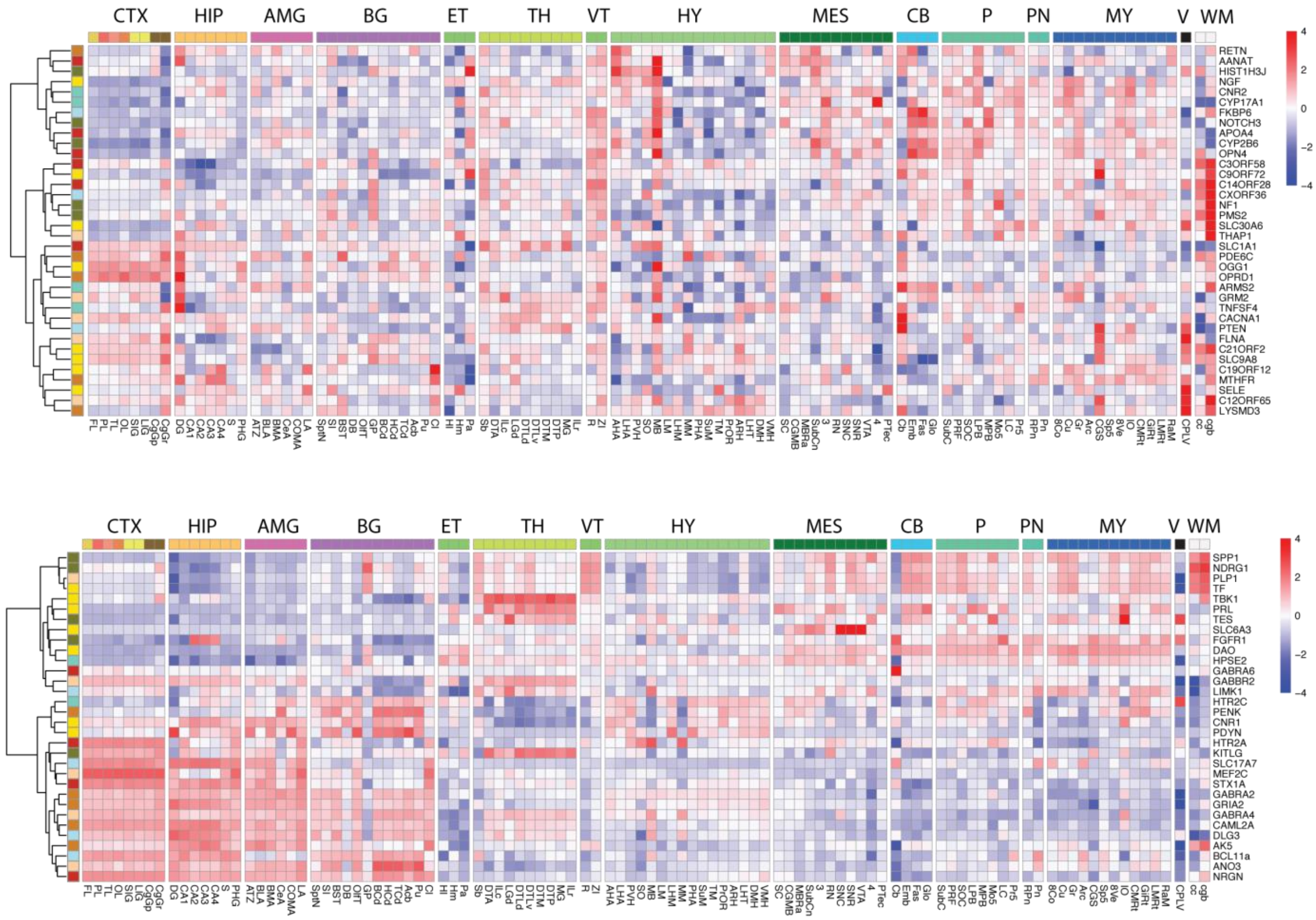


**Supplementary Figure 11. Anatomic markers for DS genes.** For each of the 40 diseases the highest and lowest differentially stable (DS) genes are selected. This results in 36 unique genes for low DS and 32 for high DS whose expression profiles are shown top (low DS), bottom (high DS). High DS genes select for structural anatomic markers and cell types. The general consistency, less random, and reduced variation is seen for the expression profile of high DS genes.

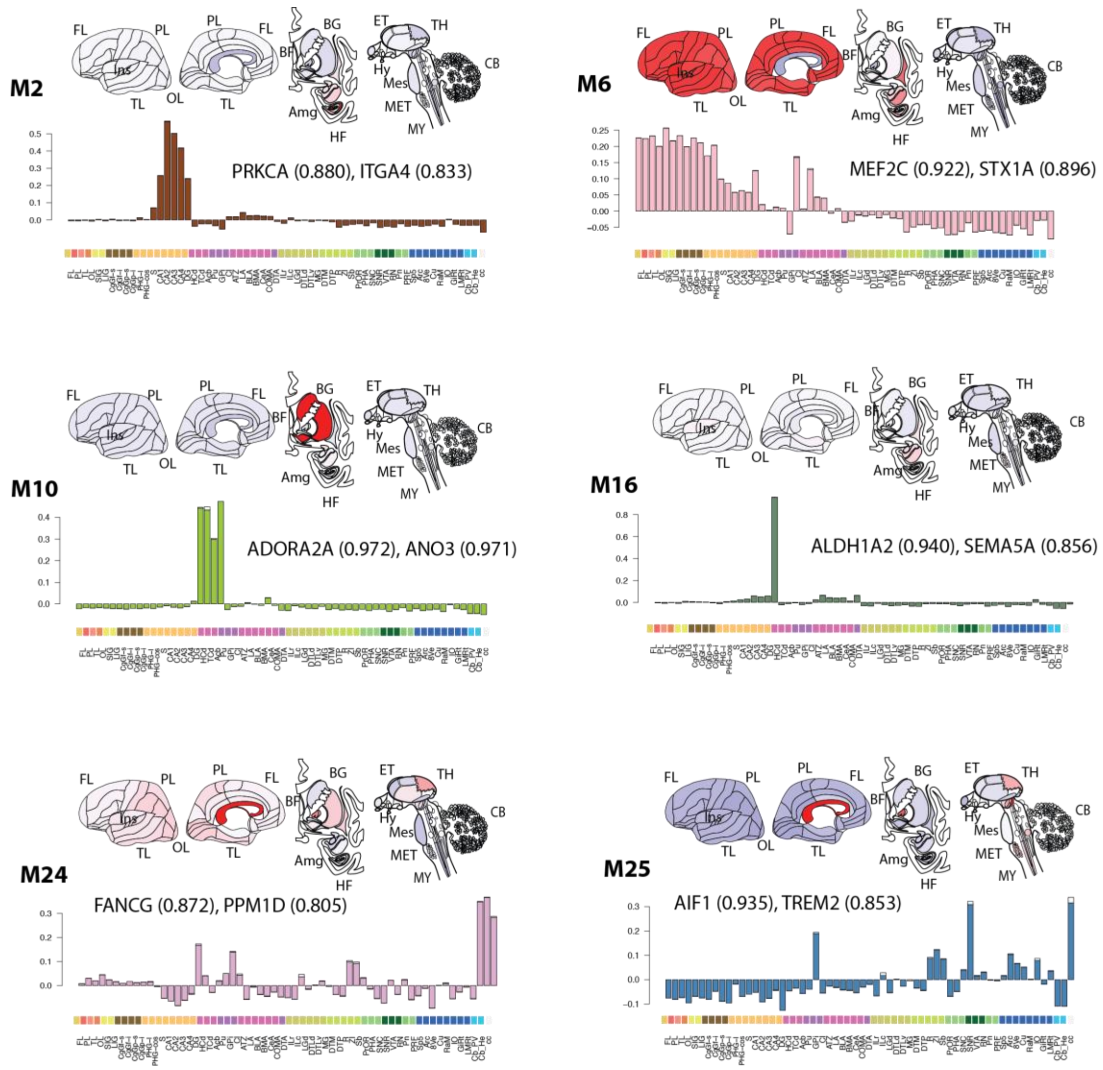


**Supplementary Figure 12. Disease associated canonical expression modules.** Canonical module M1-M32 expression patterns are highly consistent across all six AHBA individuals, and patterns identified using any five brains could be found reproducibly in the sixth (15). The modules range from structure specific markers to complex co-expression patterns in the data, and several of the modules are specific to the **ADG 1-5** groups. In addition to M1, M12 cited in the manuscript, M2 defines hippocampal expressing genes and M6 cortex-hippocampus co-expression; both are strongly represented by diseases in **ADG 3**. Representative genes and their correlation to the module eigengene are shown, PRKCA, STX1A is implicated in schizophrenia (16, 17), ITGA4, MEF2C in autistic disorder (18, 19). M10 defines striatum expressing genes and is common among ADG 3-4 diseases. ADORA2A has been studied in amphetamine-related (20) depressive disorders (21) and schizophrenia and ANO3 in dystonia (22), Parkinson's disease, ALDH1A2 in Parkinsonian disorders (23) and schizophrenia (24), SEMA5A, autistic disorder (25). Modules M24 and M25 are highly glial enriched and common in **ADG 1-2** diseases and effectively absent in ADG 3-5. FANCG has been studied in neurofibromatosis

1 (26), PPM1D in glioma (27), AIF1, Parkinson's disease (28), and TREM2 in Alzheimer's disease (29), amyotrophic lateral sclerosis (30).

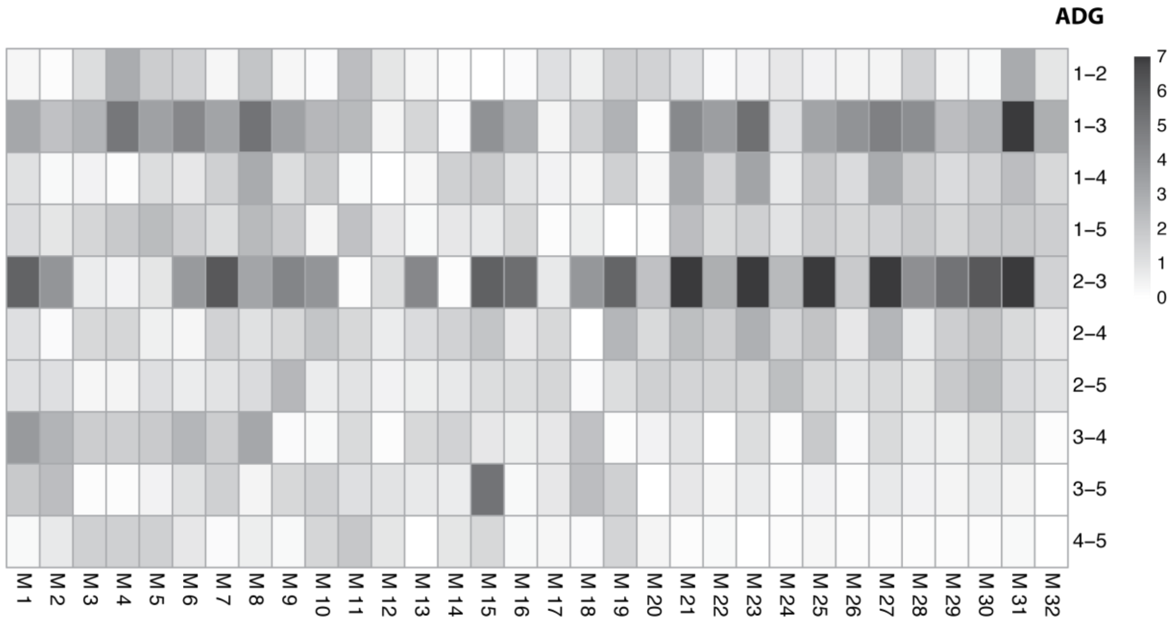


**Supplementary Figure 13. ADG group comparison within canonical modules.** Corrected t-tests between ADG groups for average disease correlation to the 32 canonical modules **M1-32.** Each set of data in the test consists of the correlation values in **Figure 2C** for those diseases in the corresponding ADG group at a fixed module. The tests are performed for all 6 pairs and each module independently. The -log10 Benjamini-Hochberg corrected values shown further validate the clustering of **Figure 1** and provide more insight into the cell patterning of ADG groups.

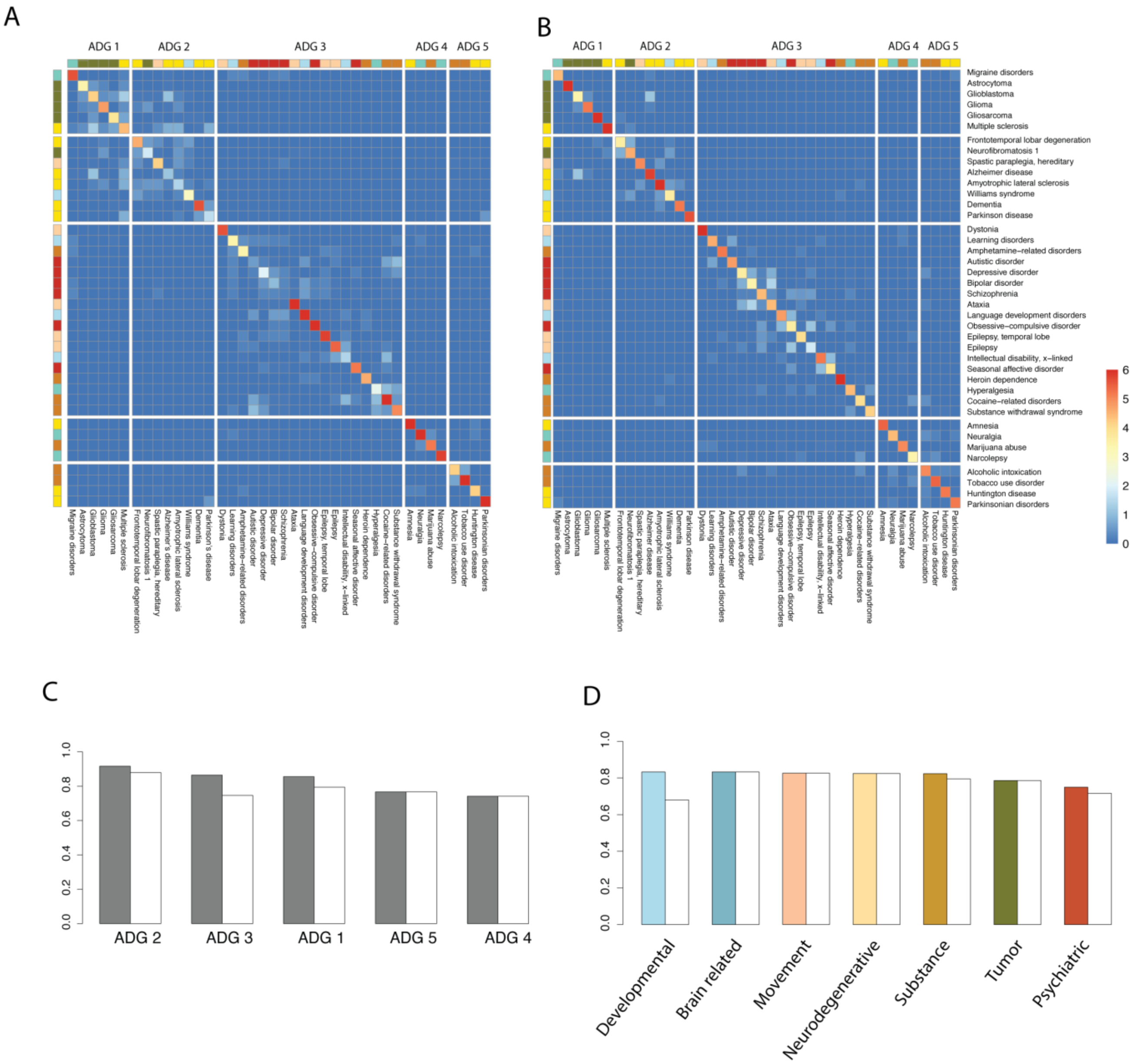


**E**

| *ADG Holdout* | ADG 1 | ADG 2 | ADG 3 | ADG 4 | ADG 5 | Mean |
|---|---|---|---|---|---|---|
| **Mean - ADG** | 0.967 | 0.771 | 0.961 | 1.0 | 0.925 | 0.924 |
| **Mean - Exact** | 0.716 | 0.537 | 0.644 | 0.875 | 0.875 | 0.746 |
| **Module - ADG** | 0.855 | 0.916 | 0.868 | 0.741 | 0.766 | 0.829 |
| **Module - Exact** | 0.794 | 0.879 | 0.746 | 0.741 | 0.766 | 0.785 |

| *GBD Holdout* | Psychiatric | Substance | Movement | Neuro-degenerative | Tumors | Develop. | Brain Related | Mean |
|---|---|---|---|---|---|---|---|---|
| **Mean - GBD** | 0.555 | 0.895 | 0.940 | 0.758 | O.753 | 0.640 | 0.825 | 0.766 |
| **Mean – Exact** | 0.433 | 0.825 | 0.933 | 0.633 | 0.606 | 0.620 | 0.816 | 0.659 |

| | | | | | | | | |
|---|---|---|---|---|---|---|---|---|
| **Module - GBD** | 0.750 | 0.823 | 0.826 | 0.825 | 0.786 | 0.833 | 0.833 | 0.811 |
| **Module - Exact** | 0.716 | 0.795 | 0.826 | 0.825 | 0.786 | 0.680 | 0.833 | 0.780 |

**Supplementary Figure 14. Holdout analysis on canonical modules and ADG.** Comparison of holdout analysis for *mean* profile of **Figure 1** and based on *canonical modules* **Figure 2**. A) Reproduction of holdout analysis for AHBA *mean* profile as in **Suppl. Figure 6** (upper diagonal.) In each of six Allen Human Brain Atlas (AHBA) subjects the mean disease transcription profile across structures is computed and the most similar (Euclidean distance) disease in the remaining 5 subjects is identified. The matrix shows the distribution of identified diseases with key 0-6 indicating the number assignments to given disease. Perfect agreement in all subjects is a 6. B) Similar analysis using *canonical module* assignments for six AHBA brains. Module based assignment shows better definition of **ADG 2** and less variance in **ADG 3** with main psychiatric diseases, bipolar, schizophrenia, autistic disorder and depression more closely identified. C,D) Classification results by ADG and GBD categories. E) Performance results for ADG and GBD comparing mean and module profiling. Mean is based on **Fig 1**, **Suppl. Fig 6** analysis; Module based on canonical module assignments. ADG or GBD label indicates that the correct class was identified, Exact indicates that precise disease was identified. Mean ADG class is reduced 10% for modules but exact disease specification is improved 4%, while for GBD groupings there is both improvement of 4.5% across all classes and for 4% exact disease identification.

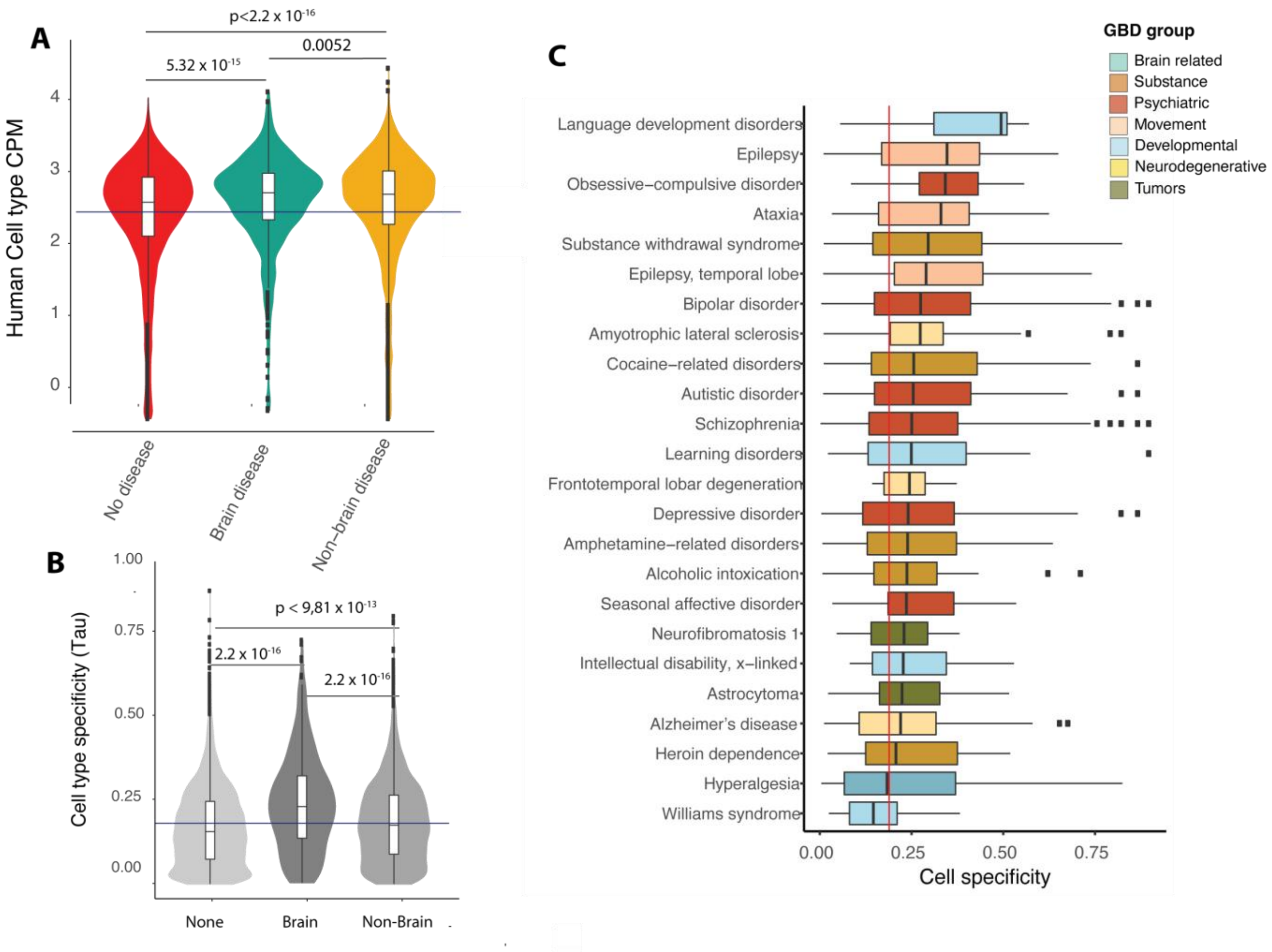


**Supplementary Figure 15. Human MTG cellular data, expression level, specificity and diseases.** A) RNA-seq gene expression quantification with absolute expression levels estimated as counts per million (CPM) using exonic reads from (31). B) Cell-type specificity was calculated based on the Tau-score (τ) defined in (32). This measure has previously been employed using the same dataset (31). Distribution of τ for brain disease associated, non-brain disease, and unassociated genes (32) C) Bar distribution plots for cell type specificity for 24 cortex expressing diseases, ordered by median specificity and colored by phenotypic GBD class. The correlation between the cell type specific tau score and the mesoscale differential stability metric is 0.445.

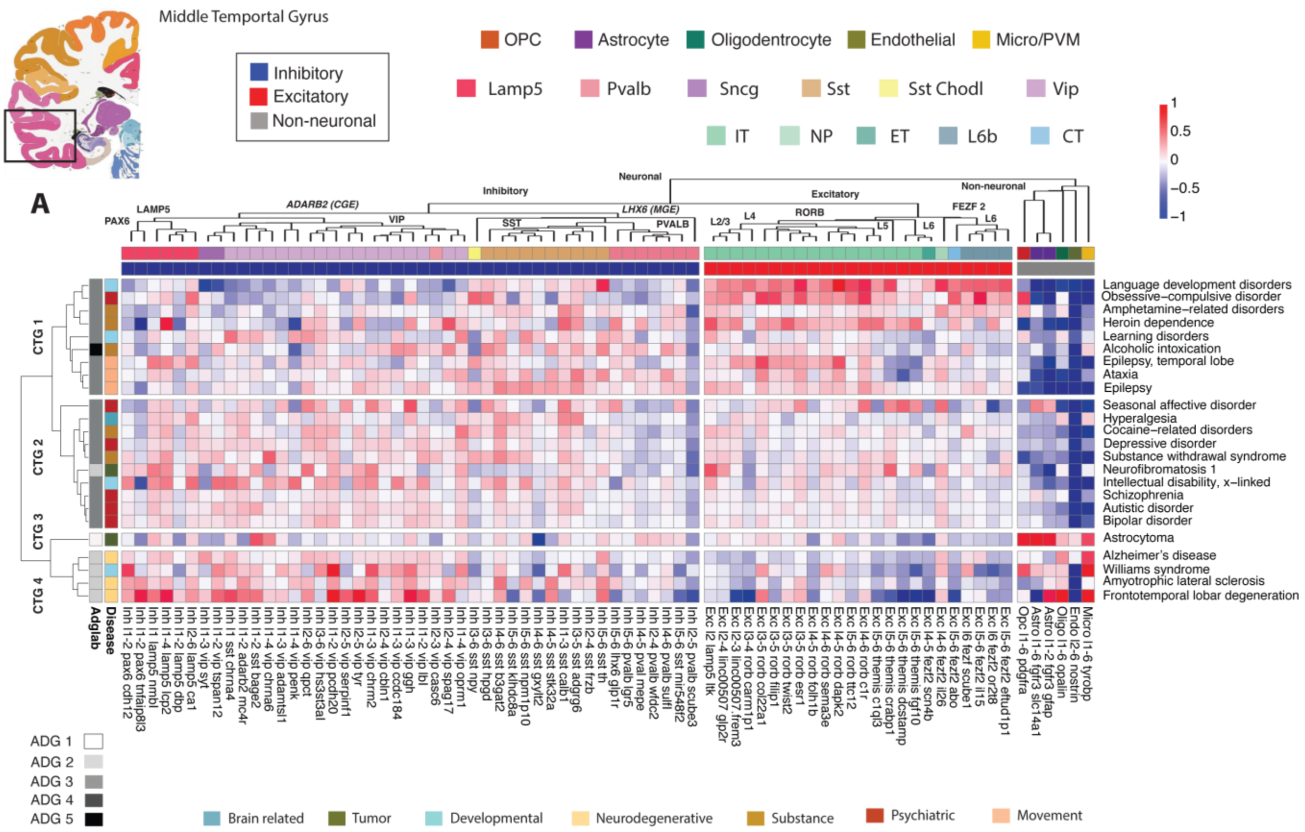

**Supplementary Figure 16. Clustered disease genes of middle temporal gyrus.** Mean cell type expression of all 40 brain diseases over 75 types identified in (31), as in **Figure 3**, but clustered only over diseases. The strong expression of excitatory types in **CTG 1** and non-neuronal types in **CTG 4**, as well as the excitatory gradient is apparent. Top annotation is reproduced from (31) and organized by major cell class; excitatory, inhibitory, non-neuronal, and color coded by subclass level; inhibitory (Lamp5, Sncg, Vip, Sst Chodl, Sst, Pvalb), excitatory (L2/3 IT, L4 IT, 5 IT, L6 IT, L6 IT Car3, L5 ET, L5/6 NP, L6 CT, L6b), non-neuronal (OPC, Astrocyte, Oligodendrocyte, Endothelial, Micro-glial/perivascular macrophages.) Left annotation: **ADG** grouping and phenotype GBD classification.

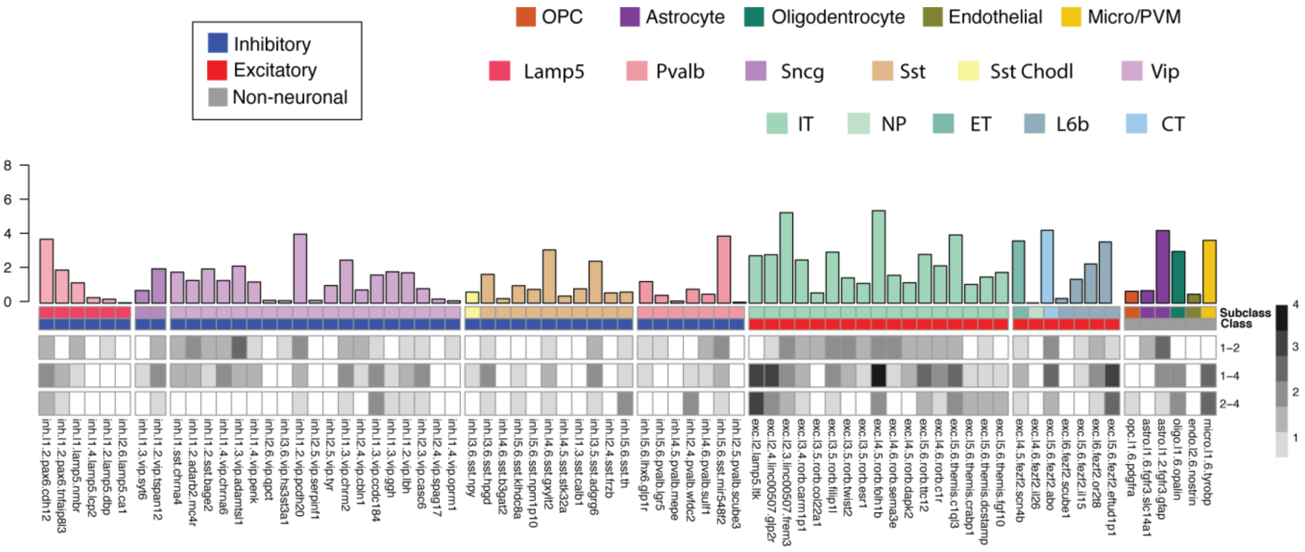


**Supplementary Figure 17. Comparing cell type clusters (CTG).** Corrected paired t-tests are used to compare significant expression differences between pairs of CTG groups, e.g. CTG 1 – CTG 2, at a fixed cell type. Overbar: ANOVA at each of 75 fixed cell types and clustered as in **Fig. 3** over three CTG groups. The highest variability is among excitatory and non-neuronal cell types and at the subclass level GABAergic Vip cell types, consistent with the excitatory and inhibitory gradients of **Fig. 3.**

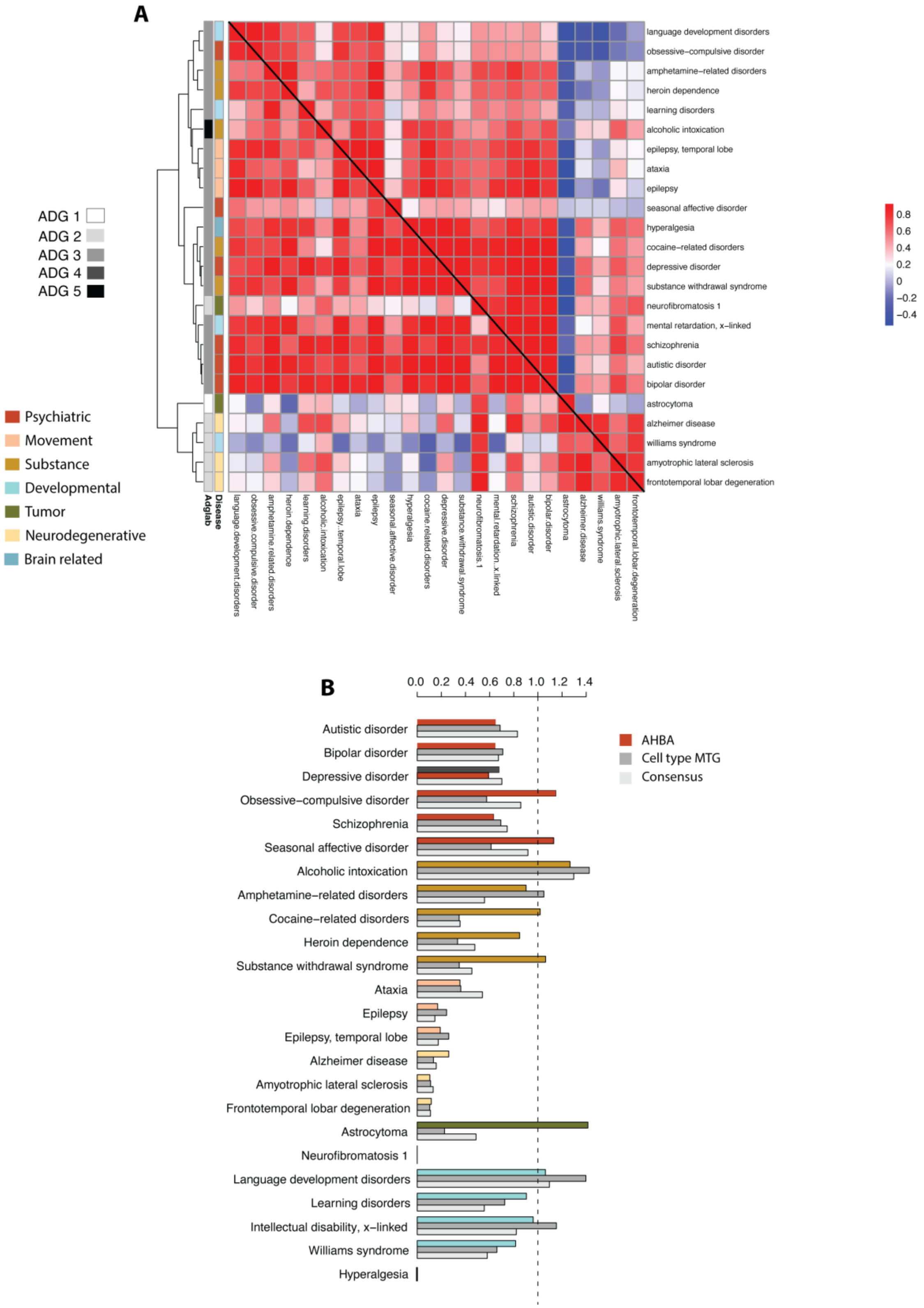


**Supplementary Figure 18.** A) Clustering matrices for correlation between 24 cortically expressing diseases based on non-overlapping genes for both HBA and cell type MTG data. Data is shown for both matrices (upper diagonal MTG, lower diagonal HBA) with clustering based on MTG data of **Figure 3**. There is general structural correspondence of these matrices and overall disease-disease Pearson correlation between the matrices is ρ=0.615. B) For each of these two-dimensional embeddings and each disease, the mean Euclidean distance from each disease to other diseases within the same GBD group is computed, as well as the mean distance to diseases not in that GBD group. The ratio of these quantities $GBD(d_i)$ is a measure of relative association of that disease with other diseases in the same GBD class. In symbols, as $GBD(d_i) = \mu_{d_j \in GBD} ||d_i - d_j|| / \mu_{d_j \notin GBD} ||d_i - d_j||$. Diseases are then grouped by their GBD class showing general agreement between the approaches, except astrocytoma which is a significant outlier better classified using the mesoscale HBA data. Solid color: AHBA brain wide, dark gray: MTG cell type, light gray: consensus.

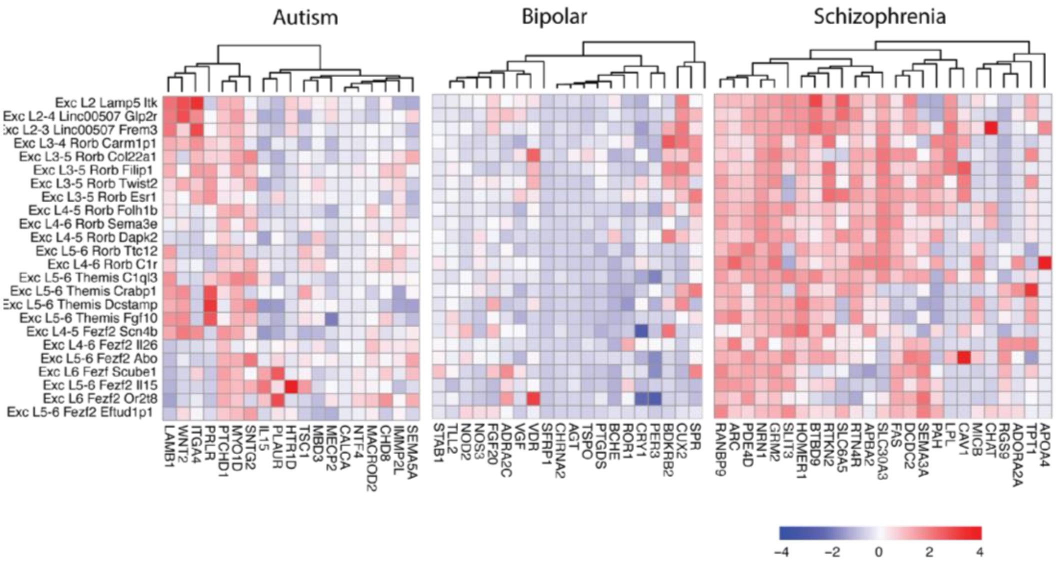


**Supplementary Figure 19. Expression profile of unique genes in autism, bipolar disorder, and schizophrenia.** Gene expression normalized for uniquely expressing genes in autism (n=19), bipolar disorder (n=20), and schizophrenia (n=25) clustered by expression level over 24 excitatory cell types. The three diseases show distinct expression profiles across excitatory types with schizophrenia widely expressing most genes.

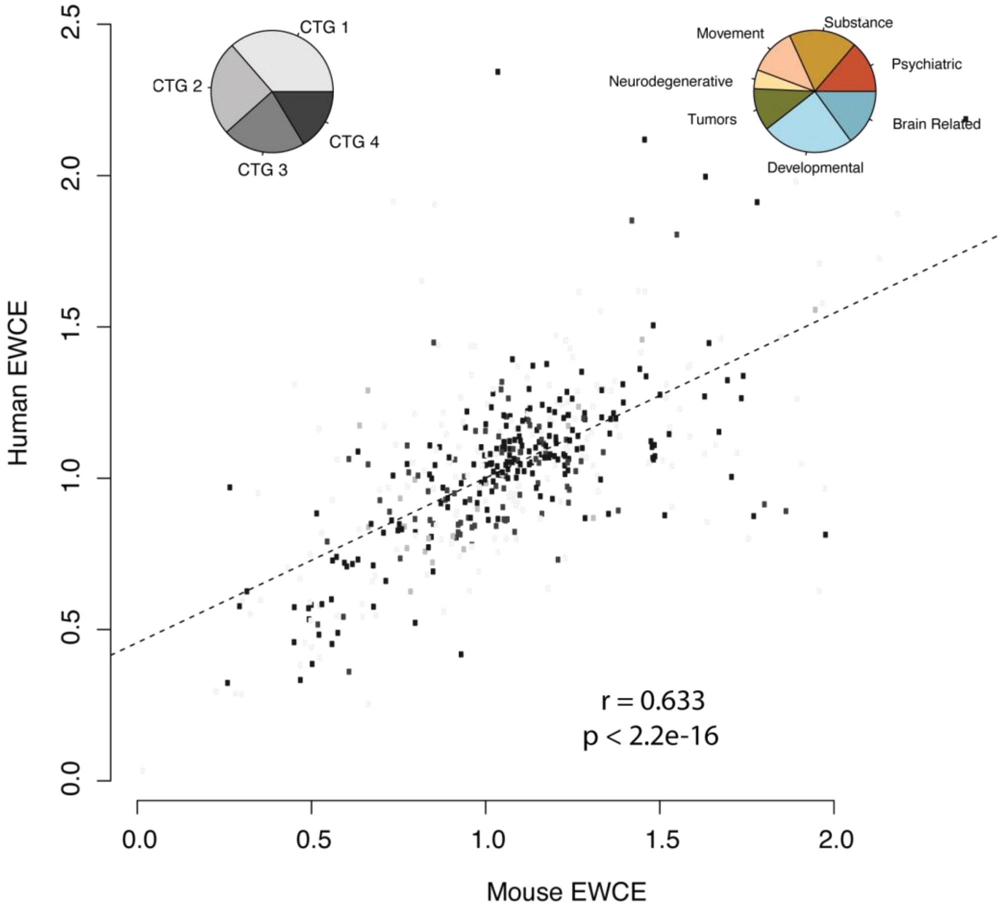


**Supplementary Figure 20. Human and mouse EWCE distributions.** Aligned transcriptomic taxonomy of cell types in human MTG to two distinct mouse cortical areas, primary visual cortex (V1) and a premotor area, the anterior lateral motor cortex (ALM) from (33) allows comparison of cell type enrichments between species. Scatterplot of disease-subclass EWCE values for mouse and human colored by **CTG 1-4.** Pie chart insets show percentages of CTG and GBD phenotypic classes of top 10% outliers from the regression line, representing most significant EWCE differences. Percentages (CTG 1, 0.363, CTG 2, 0.252, CTG 3, 0.220 CTG 4 0.163). GBD Phenotype Psychiatric, 0.137, Substance, 0.180, Movement, 0.125, Neurodegenerative 0.05, Brain tumors, 0.112, Developmental, 0.244, Brain Related, 0.150).
